# Electrical synapse molecular diversity revealed by proximity-based proteomic discovery

**DOI:** 10.1101/2024.11.22.624763

**Authors:** Jennifer Carlisle Michel, E. Anne Martin, William E. Crow, Jane S. Kissinger, Rachel M. Lukowicz-Bedford, Max Horrocks, Tess C. Branon, Alice Y. Ting, Adam C. Miller

## Abstract

Neuronal circuits are composed of synapses that are either chemical, where signals are transmitted via neurotransmitter release and reception, or electrical, where signals pass directly through interneuronal gap junction channels. While the molecular complexity that controls chemical synapse structure and function is well appreciated, the proteins of electrical synapses beyond the gap-junction-forming Connexins are not well defined. Yet, electrical synapses are expected to be molecularly complex beyond the gap junctions. Connexins are integral membrane proteins requiring vesicular transport and membrane insertion/retrieval to achieve function, homeostasis, and plasticity. Additionally, electron microscopy of neuronal gap junctions reveals neighboring electron dense regions termed the electrical synapse density (ESD). To reveal the molecular complexity of the electrical synapse proteome, we used proximity-dependent biotinylation (TurboID) linked to neural Connexins in zebrafish. Proteomic analysis of developing and mature nervous systems identifies hundreds of Connexin-associated proteins, with overlapping and distinct representation during development and adulthood. The identified protein classes span cell adhesion molecules, cytoplasmic scaffolds, vesicular trafficking, and proteins usually associated with the post synaptic density (PSD) of chemical synapses. Using circuits with stereotyped electrical and chemical synapses, we define molecular sub-synaptic compartments of ESD localizing proteins, we find molecular heterogeneity amongst electrical synapse populations, and we examine the synaptic intermingling of electrical and chemical synapse proteins. Taken together, these results reveal a new complexity of electrical synapse molecular diversity and highlight a novel overlap between chemical and electrical synapse proteomes. Moreover, human homologs of the electrical synapse proteins are associated with autism, epilepsy, and other neurological disorders, providing a novel framework towards understanding neuro-atypical states.

## Introduction

Two forms of fast synaptic transmission co-exist throughout the nervous system: chemical and electrical. While the proteomic complexity regulating neurotransmitter release and reception at chemical synapses is extensively studied^1–5^, the molecular composition of electrical synapse structure and function is not well understood. This is despite electrical transmission occurring throughout the animal kingdom^6–9^, the presence of electrical synapses during development and adulthood^10–14^, and their associations with human disorders including myopia, autism, seizure, and degeneration after neuronal injury^15–19^. Electrical synapses are clusters of tens to thousands of gap junction (GJ) channels, referred to as GJ plaques, that allow the passage of current and small molecules directly between neurons^20–22^. The identification of the GJ-channel forming proteins, Connexins in vertebrates and Innexins in invertebrates^23–27^, revealed the molecular basis for electrical transmission in synchronizing neuronal activity^28^, expanding receptive field sizes and dampening non-correlated noise^29,30^, and establishing specialized feed forward synaptic circuits^31,32^. In addition, electrical synapses are plastic^33–36^ and their ability to potentiate and depress dynamically contributes to neural computation^37–40^. Moreover, mutations in neural Connexins disrupt sensory systems including vision and smell, rhythmic systems including circadian and estrous cycles, and central systems including those controlling learning, memory, and fear responses^41–51^. Despite the breadth of knowledge indicating that electrical synapses are critical components of neural circuits throughout the brain, the molecular mechanisms by which neural Connexins localize to subcellular compartments with cell-type specificity to regulate development, plasticity, and computation remain poorly understood^52^. In order to localize to electrical synapses, Connexins are constantly on the move, trafficking on vesicles to sites of synapse formation^53^, inserting into the membrane and traversing through the GJ plaque^54^, until ultimately endocytosing and degrading in a process of constant turnover^55^. Additionally, electron microscopy of electrical synapses reveals electron dense structures within the cytoplasm, adjacent to the GJ channels^56,57^, termed the “electrical synapse density” (ESD)^58,59^. While Connexin-associated proteins within these distinct cell biological compartments are expected to regulate the structure and function of electrical transmission, molecular insight into these proteins has been hampered by a lack of methods to specifically isolate and identify the proteins of electrical synapses.

### Biotinylation of proteins in proximity to the electrical synapse *in vivo*

To identify proteins associated with electrical synapses we employed a proximity-biotinylation based approach using the evolved protein TurboID^60,61^ connected to a neural Connexin (Fig. 1A). As proof of principle, we first engineered a neural Connexin linked to TurboID and tested for functionality using cell culture. Our previous work using zebrafish genetics has identified two neural Connexins (Cx34.1/*gjd1a* and Cx35.5/*gjd2a*) and a synapse-associated cytoplasmic scaffold (Zonula Occludens 1b (ZO1b/*tjp1b*)) that are localized to and required for electrical synapse formation and function in the zebrafish central nervous system^62^. Cx34.1 and Cx35.5 are homologous to the mammalian synaptic Cx36^63^ and ZO1b is homologous to mammalian ZO1^58^ – these proteins in zebrafish and mammals are broadly localized to electrical synapses throughout the nervous system^59,63–65^. TurboID was engineered onto Cx34.1 at a position within the conserved intracellular C-terminal tail that does not disrupt known protein-protein interactions, and that when tagged with GFP allows for membrane localization in cell culture and synaptic localization *in vivo*^66^. Engineered Cx34.1-TurboID was introduced into HEK293T cells where it robustly self-biotinylated upon supplementation with exogenous biotin (Fig. S1). To determine if Cx34.1-TurboID biotinylation disrupted interaction with a known partner, mVenus was engineered onto the N-terminus of ZO1b and introduced into HEK293T cells in the absence and presence of Cx34.1-TurboID. First, we found that mVenus-ZO1b was biotinylated in the presence of Cx34.1-TurboID (Fig. S1). Second, the Cx34.1-TurboID interaction with mVenus-ZO1b was preserved in the biotinylated state (Fig. S1). We conclude that the engineered chimeric neural Connexin retains aspects of its normal cell biological interactions and can biotinylate known interacting partners.

**Figure 1.**
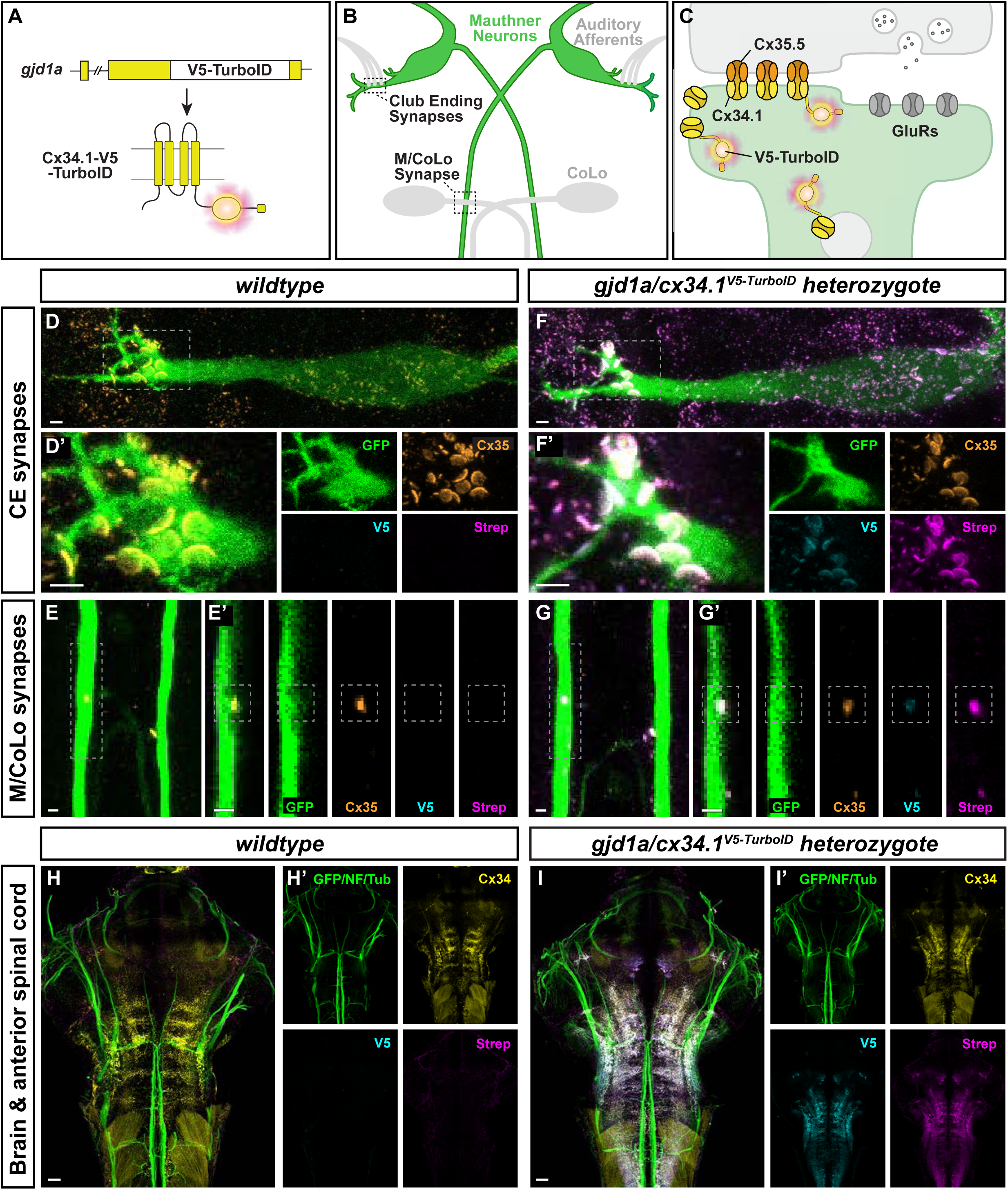
Connexin-TurboID localizes to electrical synapses and biotinylates proteins *in vivo*. **(A)** Schematic diagram of the *gjd1a/cx34.1* gene locus modified by in-frame insertion of V5-TurboID. The horizontal black bar represents the DNA strand, a white box represents V5-TurboID, and yellow boxes represent individual *gjd1a* exons. Below, a cartoon of the Cx34.1-V5-TurboID monomeric protein illustrates the insertion of the V5-TurboID cassette (magenta burst) after the four trans-membrane domains (vertical yellow cylinders) and before the C-terminal PDZ binding motif (PBM, horizontal yellow cylinder). Dark gray lines denote plasma membrane. **(B)** Simplified diagram illustrating the electrical synapses of interest in the Mauthner (M-) cell circuit. The image represents a dorsal view of the M-cells (green) with anterior on top. Regions in black dashed outline indicate the stereotypical synaptic contacts used for analysis in this study. Presynaptic auditory afferents (grey lines) contact the postsynaptic M-cell lateral dendrite in the hindbrain forming Club Ending (CE) synapses. In the spinal cord, the presynaptic Mauthner axons form *en passant* electrical synapses with the postsynaptic CoLo interneurons (grey neurons) in each spinal cord hemi-segment (1 of 30 repeating spinal segments are shown). **(C)** A diagram of a mixed electrical/chemical synapse as found at M-cell CEs. In the electrical component, molecularly asymmetric Connexin hemichannels (Cx35.5 [orange], Cx34.1 [yellow]) directly couple neurons by forming gap junction (GJ) channels. In the chemical component, presynaptic synaptic vesicles release neurotransmitter (grey circles) which align with postsynaptic glutamate receptors (GluRs). Cx34.1-V5-TurboID-containing hemichannels (yellow with magenta burst) are shown trafficking to electrical synapses on vesicles and localizing within the neural GJ plaque. **(D-G)** Enzymatically active Connexin-V5-TurboID is localized at the stereotype M-cell electrical synapses. Confocal images of M-cell circuitry and electrical synaptic contacts in 6-day-post-fertilization, *Et(T2KHG)^zf206^* zebrafish larvae from *wildtype* (D, E) and *gjd1a/cx34.1^V5-TurboID^* heterozygous (F, G) siblings treated with 1mM biotin for 72 hours. Animals are stained with anti-GFP (green), anti-Cx35.5 (orange), anti-V5 (cyan), and Strep-conjugated fluorophore (magenta). Scale bar = 2 µm in all images. Anterior up. Boxed regions denote stereotyped location of electrical synapses and regions are enlarged in neighboring panels. Images of the Mauthner cell body and lateral dendrite in the hindbrain (D, F) are maximum intensity projections of ∼27 µm. In D’ and F’, images are maximum-intensity projections of ∼15 µm and neighboring panels show the individual channels. Images of the sites of contact of M/CoLo processes in the spinal cord (E, G) are maximum-intensity projections of ∼6 µm. In E’ and G’, images are from a single 0.42 µm Z-plane and the white dashed square denotes the location of the M/CoLo site of contact. Neighboring panels show individual channels. **(H, I)** Enzymatically active Connexin-V5-TurboID is localized specifically in the brain similar to wildtype connexin. Confocal tile scans of zebrafish brain and anterior spinal cord from 6 dpf *zf206Et* zebrafish larvae from wildtype (H) or *gjd1a/cx34.1^V5-TurboID^* heterozygous (I) animals treated with 1mM biotin for 48 hours. Images are maximum intensity projections of ∼82 µm. Animals are stained with anti-GFP/anti-NF/anti-tubulin (green), anti-Cx34.1 (yellow), anti-V5 (cyan), and Strep (magenta). Neighboring panels show the individual channels. Scale bars = 20 µm. Anterior up.

To examine TurboID function at electrical synapses *in vivo*, we generated a transgenic animal expressing the Cx34.1-TurboID chimeric protein from the endogenous Cx34.1-encoding *gjd1a* gene locus. Cx34.1 is expressed in neurons found throughout the zebrafish central nervous system, including the retina, forebrain, hindbrain, and spinal cord, and is not expressed in non-neural cells^63,67^. We used CRISPR-mediated homologous recombination^68^ to introduce TurboID within the Cx34.1 intracellular C-terminal tail in the location that was successful in cell culture (Fig. 1A, S2) and developed a transgenic line of animals. To determine if the chimeric Cx34.1-TurboID protein localizes correctly in transgenics, we examined the electrical synapses of the Mauthner (M-) cell circuit. The M-cell is uniquely visualizable given its large size and distinct bipolar morphology with cell body and dendrites within the hindbrain and an axon extending down the length of the spinal cord (Fig. 1B)^63,69^. The M-cell lateral dendrites make stereotyped electrical synapses with auditory afferents of the eighth cranial nerve, termed ‘club ending’ (CE) synapses. CE synapses are ‘mixed’ electrical/chemical synapses, containing both neural GJs and glutamatergic chemical transmission (Fig. 1C)^70^. Additionally, the M-cell axon makes electrical synapses with Commissural Local (CoLo) interneurons found in each spinal-cord segment (M/CoLo synapses)(Fig. 1B). As with endogenous Cx34.1^63^, Cx34.1-TurboID localized to M-cell CE and M/CoLo synapses and colocalized with Cx35.5 at the Mauthner electrical synapses (Fig. 1D-G, Fig. S3). Additionally, Cx34.1-TurboID localized to electrical synapses throughout the larval zebrafish brain (Fig. 1H,I, Fig. S3). To examine if Cx34.1-TurboID functioned *in vivo*, we supplemented exogenous biotin into the water of larval zebrafish during a developmental window of high synaptogenesis (4-6 days post fertilization, dpf) and found extensive biotinylation of proteins at both CE and M/CoLo synaptic locations (Fig. 1D-G) as well as throughout the developing brain (Fig. 1H,I). Animals heterozygous for the Cx34.1-TurboID insertion showed greater synaptic localization and biotinylation at M-cell synapses than their homozygous counterparts (Fig. S3, Table S1), suggesting that in heterozygotes untagged Cx34.1 facilitates synaptic localization, similar to what was shown for Cx36-GFP in mouse^71^. Animals that did not have exogenous biotin added showed weaker synaptic biotinylation (Fig. S3). Given these results, subsequent work used Cx34.1-TurboID heterozygote animals supplemented with biotin to increase proximity-dependent biotinylation of Connexin-associated proteins. We conclude that Cx34.1-TurboID localizes to neural gap junctions *in vivo* and effectively biotinylates proteins found in proximity to the electrical synapse.

### Identification of neural Connexin-associated proteins

To identify neural Connexin-associated proteins *in vivo* we purified biotinylated proteins from Cx34.1-TurboID heterozygotes and wildtype controls, during both development (6 days post fertilization (dpf), whole embryos) and at adulthood (11.5 months, isolated brain), and analyzed them by high-resolution liquid chromatography-mass spectrometry (Fig. 2A). For both the developmental and adult datasets, three biological replicates of Cx34.1-TurboID or wildtype controls were prepared. Sex was not determined for developing animals as zebrafish do not have determinate sex until ∼3 weeks of age^72^. Biological replicates of female and male brains were separately prepared at adulthood. Technical replicates were performed for two of the developmental datasets. In total, spanning both the developing and adult datasets, 1974 proteins were identified, with 272 proteins enriched in the Cx34.1-TurboID samples as compared to the wildtype controls (Fig. 2B, Table S2). Developmental and adult stages had 68 and 250 enriched proteins, respectively, with 53 overlapping proteins between stages (Fig. 2B,C). Of the total enriched proteins, 29 and 146 proteins in the developing and adult datasets, respectively, were significantly enriched based on a strict false discovery rate (Table S2). Additional candidate proteins considered lower-confidence, yet absent from wildtype controls, were consistently present at greater than 7-fold enrichment in all replicates of developing or adult Turbo-ID data (Fig. 2C, Table S2). No differences were observed in enriched proteins within technical replicates of the developing datasets nor between female and male adult brains (Table S2).

**Figure 2.**
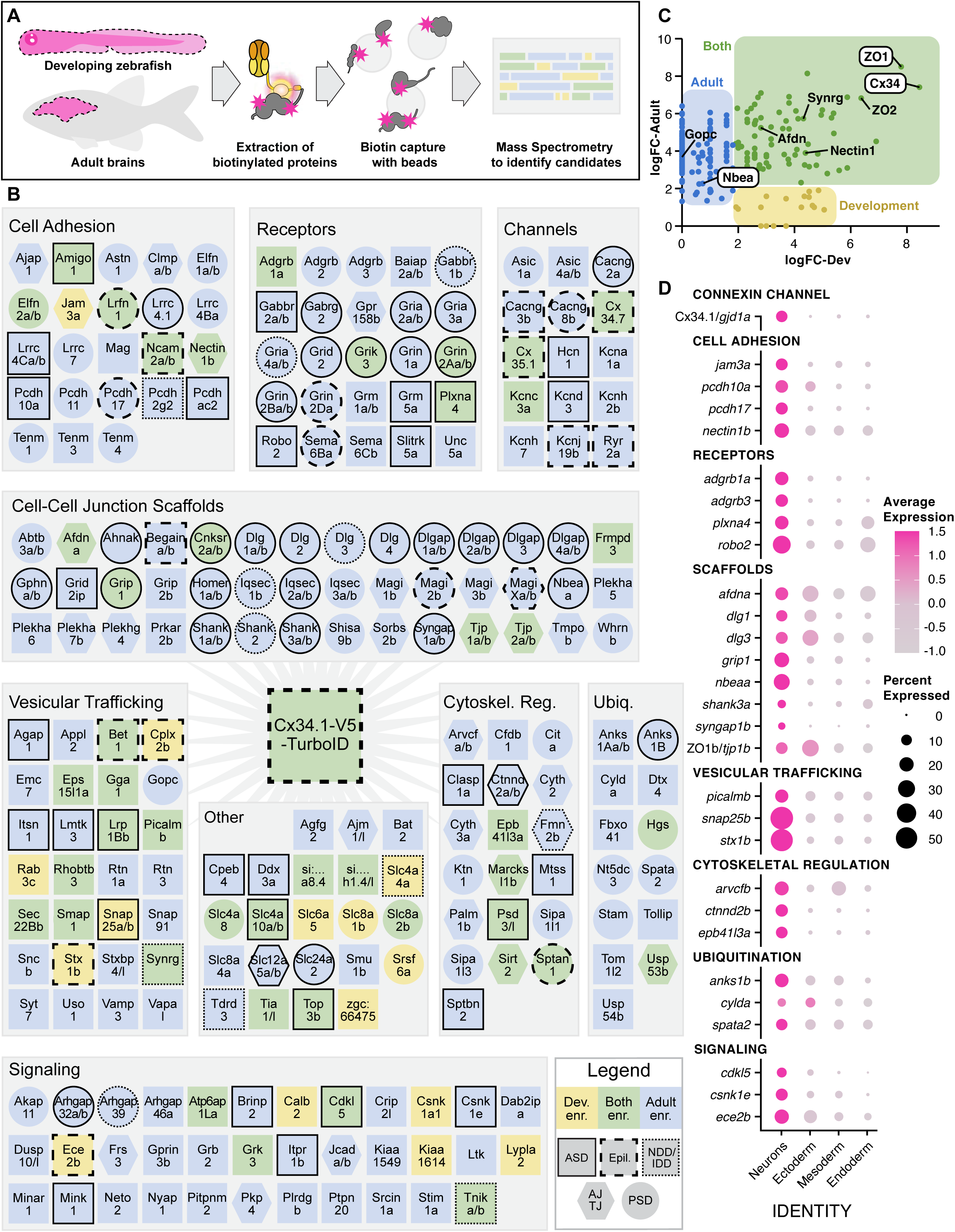
Discovery of the neural Connexin-associated proteome by biotin-proximity identification. **(A)** Diagram of biotin-proximity labeling scheme. *gjd1a/cx34.1^V5-TurboID^* heterozygous larvae (developing) or adults were treated with biotin supplemented water to activate maximal biotinylation of nearby proteins. Whole larvae and brains from adults (dotted lines) were harvested and homogenized under denaturing conditions. *gjd1a/cx34.1* is expressed exclusively in the nervous system providing specificity. Biotinylated proteins were captured with magnetic streptavidin beads, digested with trypsin on-bead, and analyzed by mass spectrometry to identify candidates. **(B)** Proteins enriched in Cx34.1-V5-TurboID samples compared to wildtype controls are depicted and grouped into grey boxes according to gene ontology (GO) analysis: Cell Adhesion, Receptors, Channels, Cell-Cell-Junction Scaffolds, Vesicular Trafficking, Signaling, Cytoskeletal Regulation, Ubiquitination, and Other. Paralogous proteins are denoted as single shapes and noted by a forward slash between name delineations. Colors denote enrichment in the developmental (yellow), adult (blue), or both datasets (green). Hexagons denote proteins associated with epithelial adherens or tight junctions (AJ/TJ), circles denote proteins associated with postsynaptic densities of chemical synapses (PSD), and squares denote the remainder of the isolated candidates. Outlines denote human disorders associated with the protein: solid outlines denote proteins associated with autism spectrum disorder (ASD), dashed outlines denote proteins associated epilepsy (Epil.), and stippled outlines denote proteins associated with Neurodevelopmental Disorders and/or Intellectual Disability (NDD/IDD). Many proteins have associations with multiple of these categories, or other neurological disorders (Table S2). **(C)** Enrichment of identified proteins (dots) in developing (x-axis) and adult (y-axis) datasets. Colored regions denote proteins enriched in developing (yellow), adult (blue) or both (green) time points. Proteins in white callouts have been previously localized to zebrafish electrical synapses (Cx34.1, Nbea, and ZO1) while the rest have been localized to mammalian electrical synapses or interact with mammalian Cx36. **(D)** Neural Connexin-associated proteome gene expression across germ layers in a whole-embryo single-cell RNA sequencing dataset (5 and 6 dpf). Clusters (Identity) are organized by germ layer along the x-axis: neurons, ectoderm (non-neural ectoderm, e.g. skin and cranial neural crest), mesoderm (e.g., skeletal muscle and blood), and endoderm (e.g. liver). A selection of Connexin-associated genes is arranged along the y-axis, grouped by GO terms. Dot size represents the percentage of cells in the identity category expressing each gene, while color indicates relative expression levels.

To assess the quality of enrichment in the dataset, we first cross referenced the identified proteins with gene expression captured in our single cell RNA-seq atlas of zebrafish development^67,73^. We found that enriched candidates were biased towards mRNA expression in neurons, with notable co-expression with endogenous Cx34.1/*gjd1a* (Fig. 2D, S4). Given the partial genome duplication of teleost lineages^74^, we used BLAST and the Zebrafish Information Network (ZFIN)^75^ to identify 216 human proteins with clear 1-to-1 orthology in the enriched protein dataset, with only 4 proteins absent from mammalian lineages (Table S2). Gene Ontology (GO) analysis^76,77^ of the proteins highlighted significantly enriched terms including postsynapse (p=6.19E-67), neuron-to-neuron synapse (p=1.01E-65), dendrite (p=2.27E-35), and trans-synaptic signaling (p=9.80E-33) (Table S2), suggesting we had enriched for proteins localized at neuronal synapses. In line with this enrichment, GO analysis, cross referenced with resources cataloging human-disease-associated genes^78–81^, revealed an abundance of proteins associated with autism spectrum disorders (ASD), epilepsy, and neurodevelopmental disorders and/or intellectual disability (Fig. 2B, Table S2). While electrical synapses are abundant throughout the brain, there are currently only ∼10 proteins known to localize to vertebrate neural gap junctions^52,82^. Within the enriched proteins, many of these known proteins that localize to electrical synapses or interact with neural Connexins were identified (Fig. 2C). The two most enriched proteins in the datasets were Cx34.1, onto which we engineered TurboID, and ZO1b, which we have previously shown is localized to the electrical synapse and binds directly to Cx34.1 via a PDZ-PBM mediated interaction^58,59,83^. In addition, Neurobeachin (Nbea) was enriched in the adult dataset, and our previous work showed it is localized to electrical synapses and required for synaptic localization of both Cx34.1 and ZO1b^66,84^. Additionally, a number of proteins previously determined to localize to mammalian electrical synapses were enriched in the datasets, including ZO2^85^, Afadin (Afdn)^86^, and Nectin1^87^(Fig. 2C). Further, the experiment is expected to enrich for proteins within proximity of Cx34.1-TurboID from translation to degradation, therefore the candidates are likely to include proteins that are not exclusively at the synapse. Indeed, also enriched in the data are the trafficking proteins Golgi Associated PDZ And Coiled-Coil Motif Containing (Gopc) and Synergin (Synrg) that were identified to be in proximity with Cx36 when it is overexpressed in HEK293T^88^. We conclude that the biotin-proximity labeling approach enriched for neuronal and synaptic proteins and successfully identified Connexin-associated proteins *in vivo*.

Analysis of known functions of the other, enriched, candidate Connexin-associated proteins, or of human and mouse orthologs (Table S2)^76^, highlighted a number of putative functions that have been hypothesized to regulate electrical synapse assembly and structure^52,53,89^. These included neural cell adhesion molecules, neural signaling and synaptic receptors, cytoplasmic scaffolds, and cytoskeletal proteins (Fig. 2B). Additionally, Connexins are four-pass transmembrane domain proteins that oligomerize into hemichannels in the ER/Golgi, are trafficked in vesicles to sites of GJ formation, and are removed and degraded by the proteosome^20,90,91^. The dataset identified ER and Golgi proteins, vesicular trafficking regulators, and ubiquitination pathway proteins (Fig. 2B) that may regulate neural Connexins. Two additional major cell biological categories emerged from the analysis: (1) epithelial adherens and tight junction (AJ/TJ) proteins (Fig. 2B, hexagons) and (2) chemical synapse postsynaptic density (PSD) proteins (Fig. 2B, circles). Recent work has revealed the close association of a subset of AJ/TJ proteins with neural Connexins^52,82^. Regarding the ‘chemical synapse PSD’ proteins, it is noteworthy that these candidate proteins have not been studied in the context of the electrical synapse, raising the possibility that they may also be ESD proteins. Additionally, zebrafish contain an abundance of ‘mixed’ synapses wherein individual neuronal contacts exhibit simultaneous electrical and chemical transmission^92,93^. Mixed synapses are also found in mammalian and invertebrate neural circuits^56,94^, though are not well studied. Taken together, we conclude that the approach identified known neural Connexin-associated proteins and highlights further candidate molecules with a diversity of putative cell biological functions.

### Localization of Electrical Synapse Density proteins *in vivo*

To identify proteins localized to synaptic contacts, the ESD proteins, we investigated the localization of candidate Connexin-associated proteins *in vivo*. We endogenously tagged candidate proteins from amongst the major identified functional classes and examined their localization in developing zebrafish. Given the general lack of antibodies available for zebrafish proteins, we used CRISPR to introduce V5-tags into proteins at their endogenous genomic loci^95^ and used immunofluorescence to examine the localization of the tagged proteins in relation to Cx34.1 (Fig. 3A). First, we examined the brain-wide expression patterns of candidate proteins and found a diversity of patterns, including expression throughout the brain, those with regionalized expression, including anterior (forebrain/midbrain) or posterior biases (hindbrain/spinal cord), or patterns with sparse and cell-type specific expression in subsets of neurons (Fig. 3B). In addition, protein localization was found in a diversity of other tissue types, including glia, epithelia, muscle, and vasculature (Fig. S5). Given that the approach produces mosaic embryos^66,95^, we examined multiple examples of each investigated protein and found consistent patterns of expression and localization (Fig. S5). Throughout the diversity of neural expression classes, regions of overlapping expression with Cx34.1 were identified for a majority of proteins examined (85%, 33/39) (Fig. 3, S5). Cases lacking overlap could be due to an inability to visualize the protein at locations of colocalization, that the proteins do not colocalize at this time of development, or that V5 tagging disrupted endogenous localization. For those with Cx34.1 overlapping localization, some proteins showed colocalization throughout the brain while others displayed only regional colocalization (Fig. 3B), suggesting that electrical synapses at distinct locations have unique proteomic makeups. From these broad categorizations of Connexin-associated proteins it appears that the candidate proteins examined show a high degree of specific neuronal expression, frequently colocalize with Cx34.1, and that distinct populations of electrical synapses may have unique molecular constituencies.

**Figure 3.**
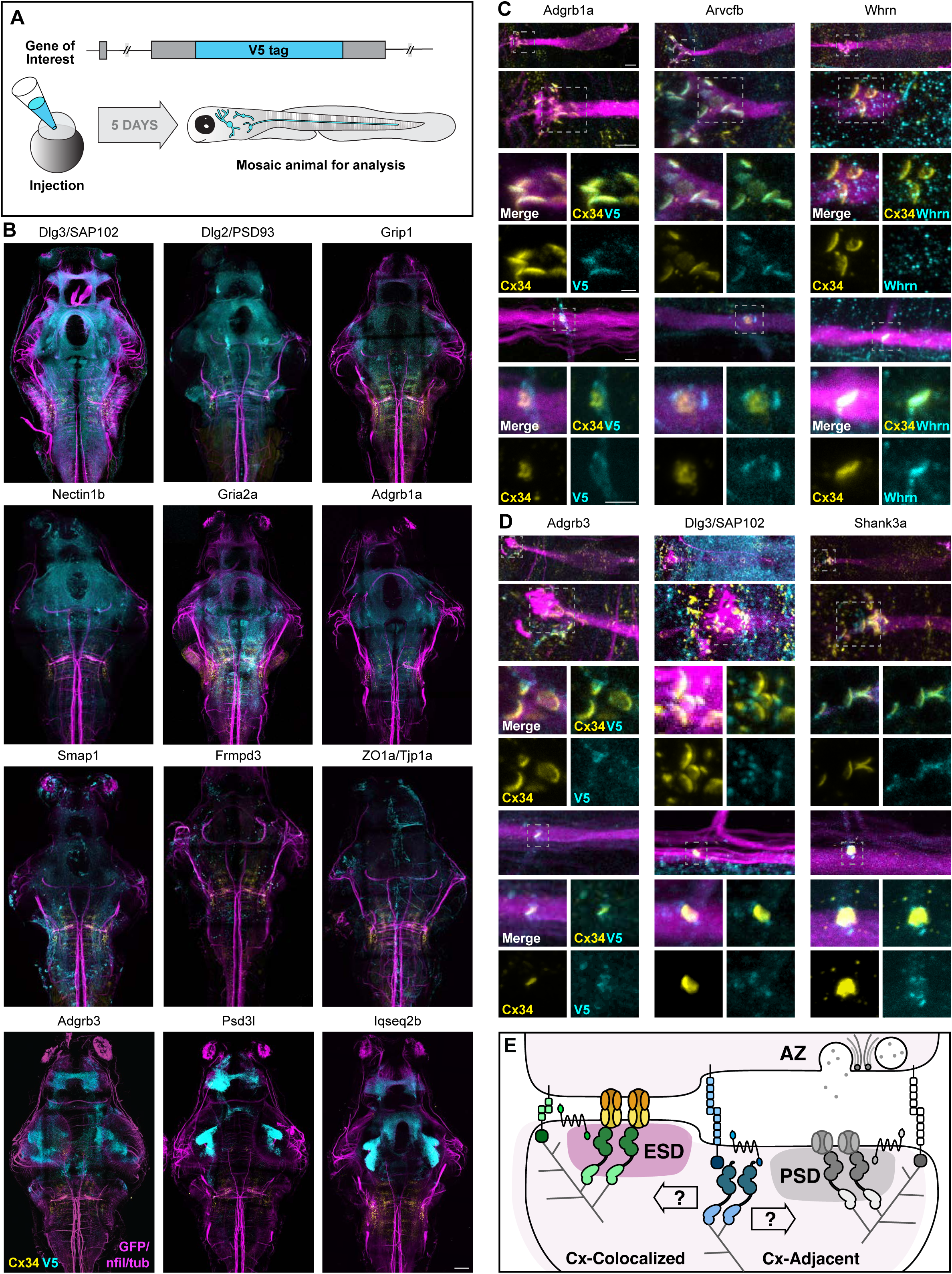
Identification of electrical synapse density proteins. **(A)** Diagram of CRISPR-mediated epitope insertion at endogenous genes to generate animals mosaically expressing V5-tagged proteins for analysis in the zebrafish brain. The horizontal black bar represents the DNA strand of the gene of interest, a cyan box represents the insertion of the V5 epitope tag. Animals are injected at the 1-cell stage and expression is visualized five days later. **(B)** Confocal tile scans of zebrafish brain at 5 dpf *Et(T2KHG)^zf206^* zebrafish larvae expressing V5-tagged proteins, as labeled. Animals are stained with anti-GFP/anti-NF/anti-tubulin (magenta), anti-Cx34.1 (yellow), and anti-V5 (cyan). Scale bars = 20 µm. Anterior up. **(C)** Confocal images of Connexin-colocalized genes in the M-cell circuit at 5 dpf in *Et(T2KHG)^zf206^* zebrafish larvae. Boxed regions denote stereotyped location of electrical synapses and regions are enlarged in neighboring panels. Animals are stained with anti-GFP/anti-NF/anti-tubulin (magenta), anti-Cx34.1 (yellow), and anti-V5 or anti-Whrn antibody (cyan), as labeled. Top four rows show CE synapses, anterior up. Bottom three rows show M/CoLo synapses, anterior left. Neighboring panels show individual and merged channels, as labeled. Scale bars = 2 µm. **(D)** Confocal images of Connexin-adjacent genes in the M-cell circuit at 5 dpf in *Et(T2KHG)^zf206^* zebrafish larvae. Boxed regions denote stereotyped location of electrical synapses and regions are enlarged in neighboring panels. Animals are stained with anti-GFP/anti-NF/anti-tubulin (magenta), anti-Cx34.1 (yellow), and anti-V5 (cyan). Top four rows show CE synapses, anterior up. Bottom three rows show M/CoLo synapses, anterior left. Neighboring panels show individual and merged channels, as labeled. Scale bars = 2 µm. **(E)** Model of a mixed electrical/chemical synapse illustrating Connexin-colocalized and Connexin-adjacent proteins. ESD proteins fall into classes such as cell adhesion, signaling, scaffolding, cytoskeletal regulation and colocalize with electrical synapses (green shapes). These proteins classes are analogous to proteins identified by proteomics at chemical synapse presynaptic active zones (AZ) and PSDs (grey shapes). Candidates that are adjacent to electrical synapses (blue shapes) may be exclusive to the chemical synapse, may be localized adjacent to both, and may regulate both electrical and chemical synapses assembly and function, as indicated by the opposing arrows. ESD, electrical synapse density. PSD, post-synaptic density. AZ, pre-synaptic active zone.

To investigate the subcellular localization of identified proteins we turned our attention to the M-cell circuit with its stereotypical cell biology and synapses. The M-cell CE synapses (Fig. 1B) are mixed electrical/glutamatergic chemical synaptic contacts, with EM^70,96^ and expansion microscopy^97^ showing that the chemical component of these mixed synapses residing in distinct locations surrounding the synaptic GJ plaques (Fig. S6). At M/CoLo synapses, transmission has a clear electrical component, though electrophysiology was indeterminant in relation to a chemical component at these synapses^98^. At both CE and M/CoLo synapses, Cx34.1 and AMPA receptors are localized to distinct subsynaptic domains where Connexin adopts a stereotypical ‘C-shaped’ pattern that stretches across the synaptic contact, whereas AMPA receptors are localized at 2-4 distinct puncta surrounding Cx34.1 localization (Fig. S6). Using the unique cell biological accessibility of the M-cell circuit, we investigated the localization of the V5-tagged Connexin-associated proteins expressed in the hindbrain and spinal cord where the circuit resides. Two major classes of specific, synaptic localization were identified: Connexin-colocalizing and Connexin-adjacent.

The Connexin-colocalizing protein class was typified by localization in a stereotypical C-shaped pattern at CE or M/CoLo synaptic contacts and extensively overlapped with Cx34.1 staining (Fig. 3C, S5). In total, 16 proteins were found to be Connexin-colocalizing: Adgrb1a, Afdna, Arvcfa, Arvcfb, Clmpb, Ctnnd2a, Ctnnd2b, Dlg1b, Jam3a, Nectin1b, Plekha5, ZO1a, ZO1b, ZO2a, ZO2b, and Whrnb. For Whrnb we obtained an antibody against the zebrafish protein^99^. In zebrafish, only ZO1b had previously been co-localized with neural Connexins^58,59,83^; in mouse, Afdn, Nectin1, ZO1, and ZO2 have been co-localized with Cx36^64,85–87,96^. The rest of the molecules are novel ESD proteins. It is notable that these proteins are largely, though not exclusively, associated with epithelial AJs/TJs, with little known about their function in the nervous system. The Connexin-colocalizing class represents cell adhesion molecules, cytoskeletal regulators, and cytoplasmic scaffolds (Fig. 2B). We have previously shown that ZO1b localizes to electrical synapses (Fig. S6) where it directly interacts with Cx34.1 and is required for synaptic localization and electrical transmission^58,59,83^, suggesting that this group of proteins may represent a signaling complex for the assembly, structure, and function of the electrical synapse.

The second synaptic class, Connexin-adjacent, contained eight proteins and included the glutamatergic neurotransmitter receptor subunit Gria2a, the receptor Adgrb3, and the cytoplasmic scaffolds Frmpd3, Grip1, Nbeaa, Sap102, Shank3a, and Syngap1b (Fig. 3D). Note that given our reliance on V5-tagging, and the lack of antibodies available that recognize these zebrafish proteins, the distribution of these proteins relative to one another at individual synapses cannot currently be determined. The Connexin-adjacent class of molecules is dominated by associations with chemical synapse PSD proteins (Fig. 2B). Yet, we have previously found that Nbeaa localizes in a Connexin-adjacent pattern at M-cell electrical synapses (Fig. S6), and despite this ‘next-to’ localization, it is required for robust synaptic Cx34.1 localization and M-cell induced escape response behavior^66,84^. This suggests the provocative possibility that the Connexin-adjacent class contains at least a subset of proteins with functions in electrical synapse assembly and function.

Finally, we observed that the M-cell electrical synapses displayed molecular heterogeneity. The M-cell circuit receives sensory input from auditory afferents at the CEs and transmits this information to motor circuits in the spinal cord including to the CoLo interneurons. Both Adgrb1a and Arvcfb were localized in a Connexin-colocalizing pattern at CEs in the hindbrain, but at M/CoLo synapses in the spinal cord they were localized in unique synaptic-contact-surrounding patterns (Fig. 3C, S3). This pattern appeared distinct from the Connexin-adjacent class described above. By contrast, Whrnb was found localized in a Connexin-colocalizing pattern at M/CoLo synapses but at CE contacts was localized in a Connexin-adjacent fashion. It is striking that within the M-cell escape circuit, distinct electrical synapses display molecular diversity with subcellular specificity, potentially related to the distinct functions (sensory integration at CEs, motor coordination at M/CoLos) of each synapse population.

Taken together, the synaptic proteins represent at least two distinct synaptic localization patterns, Connexin-colocalized or Connexin-adjacent. Those that are Connexin colocalized represent putative ESD proteins for which genetic, molecular, and cell biological analysis may reveal functions in electrical synapse structure and function (Fig. 3E, green). Those that are Connexin-adjacent represent putative PSD proteins, for which many are expected to have functions at the glutamatergic synapses found at Mauthner cell mixed synapses (Fig. 3E, gray). Yet the Connexin-adjacent class also highlights the emerging possibility of molecular and functional overlap between the ESD and PSD proteins (Fig. 3E, blue).

## Discussion

The Connexin-associated proteome presented here reveals hundreds of new molecules we hypothesize can fulfill the varied and rich roles of electrical synapse regulation during assembly and function. These include proteins that fall into classes such as cell adhesion, signaling, scaffolding, cytoskeletal regulation, and vesicular trafficking – all categories analogous to those identified by proteomics at chemical synapse presynaptic active zones (AZ) and PSDs (Fig. 3E)^100–105^. Within the ∼250 proteins identified here we localized sixteen proteins to Connexin-colocalized and eight to Connexin-adjacent locations at M-cell electrical synapses and observed similar distributions at other Cx34.1 puncta found throughout the brain. Localization of proteins also revealed molecular heterogeneity at M-cell synapses, highlighting that not all electrical synapses are made equal, but instead that they can have unique molecular compositions likely to affect neural gap junction structure and transmission. Additionally, the proteomic data highlights proteins that track the expected ‘life cycle’ of Connexins, from vesicular trafficking in the ER/Golgi, to synaptic associations between neurons and with cytoplasmic regulators, to protein degradation after endocytosis^20^. While only a subset of these molecules will be strictly ‘synaptic’, therefore being defined as ESD proteins, the entirety of the identified proteome may regulate neural Connexin biology. A key frontier that emerges is to define the electrical synapse beyond the Connexins and to understand the molecular framework that builds its structure. Critically, we must convert the gross synaptic localization patterns observed here into the sub-synaptic localization at the electrical synapse. EM of electrical synapses highlights the GJ plaques containing hundreds to thousands of channels, and also reveals the directly adjacent puncta adherens, and the cytoskeletal complexity^12,56,106^. The molecular richness of the electrical synapse proteome is expected to define synaptic domains that regulate the structure, transmission, and plasticity of electrical transmission.

The neural Connexin-associated data here is expected to reveal only a subset of the complete electrical synapse proteome. First, proteins that are not within proximity of the Cx34.1-TurboID chimeric enzyme, or that are in compartments inaccessible to the enzyme, are not biotinylated, and therefore are not captured in the approach. In line with the expectation, N-cadherin (Cdh2) and Beta-catenin (Ctnnb2) were recently localized to M-cell electrical synapses^97^ but were not identified in our data. Second, more complexity is expected as we look to other electrical synapses. Here we focused on a zebrafish Cx36 homologue in fish^63^, yet there are five distinct Connexins that make electrical synapses in mammals^21,26^, and future experiments will reveal the distinct, and overlapping, proteomic compositions of the other neural Connexins. Finally, electrical synapse complexity also appears to be related to age as our data revealed differences in developing and adult animals. Such electrical synapse molecular compositions shifts could affect transmission and neural circuit function, similar to what has been observed at chemical synapses^107–109^. The data here reveals a molecularly rich proteome for electrical synapses, analogous to what is known for chemical synapses, with more complexity yet to be discovered.

A notable revelation from the Connexin-associated proteomic data was the identification of chemical synapse PSD molecules, including glutamate receptors, scaffolds, cell adhesion molecules, and regulators. These observations force us to consider where the boundaries lie between different synapse types (Fig. 3E). Electrical and excitatory chemical co-transmission is a common synaptic motif that can be found at mixed synapses between two neurons, as found in the M-cell circuit and other locations, or in cases where separate electrical and glutamatergic synapses are made between two presynaptic neurons and a single postsynaptic partner^11,92,94^. In both cases, postsynaptic electrical and chemical components could co-mingle in close proximity, or in trafficking pathways, and this could explain the observed PSD proteins in the Connexin-TurboID data. Alternatively, so-called PSD proteins may not be as exclusive as the name implies, and perhaps they also function in the regulation of electrical synapses. Notably, this proteomic work identified Neurobeachin (Nbea), which is linked to autism and epilepsy^110–112^ and is required for the localization of both electrical and chemical synapse proteins^66,84,113–116^. The notion of synapse coordination marks a frontier where we must define the proteins that belong exclusively to the electrical or chemical synapse and those that ‘moonlight’ and perform essential functions at both. Future imaging with higher spatial resolution (e.g. immuno-EM and expansion microscopy) combined with genetic, biochemical, and functional work, will reveal the mechanistic functions of synapse specific and synapse coordinating proteins.

Broadening our molecular understanding of the electrical synapse creates fundamental new insights into the complexity of neural circuits and new possibilities related to human neurological disease states. In line with this notion, neural Connexin mutants in mouse and zebrafish have revealed complex behavioral defects related to vision, smell, circadian rhythms, estrous cycles, learning and memory, fear responses, and motor coordination^41–51,63^. Additionally, human mutations affecting Cx36/*GJD2* are a leading cause of myopia^15^ and have been proposed to underlie autism^16^, while hyperfunction of electrical synapses may lead to seizure^17,18^. Here, the Connexin-associated proteome identified a myriad of proteins in which the human orthologues are linked to a variety of neurological disorders, with particular enrichment for autism, epilepsy, and neurodevelopmental and intellectual disabilities. Future experiments to reveal the mechanisms by which the Connexin-associated proteins regulate electrical synapse assembly, structure, plasticity, and coordination with chemical synapses, highlights an enticing new direction to understand human neural circuit disorders.

## Acknowledgments

Research was supported by NIH grants R21NS117967 and R01NS105758 to ACM, by the Eunice Kennedy Shriver National Institute of Child Health and Human Development of the National Institute of Health under Award Number F32HD102182 to EAM, Dow Graduate Research and Lester Wolfe Fellowships to T.C.B., and the NIH National Institute of General Medical Sciences (NIGMS) Graduate Training in Genetics Grant T32GM007413 to MH, RML, and WEC. Mass spectrometric analysis was performed by the OHSU Proteomics Shared Resource by Kilsun Kim (sample preparation), Ashok Reddy (data collection), and Phillip Wilmarth (data analysis), with partial support from NIH core grants P30EY010572, P30CA069533 and OHSU Emerging Technology Fund. We thank the University of Oregon zebrafish facility staff for superb fish care, especially through the challenges of the global pandemic. We thank the administrative staff at the University of Oregon Institute of Neuroscience. We thank Ava Komons and the entire Miller lab for critical contributions and feedback on the manuscript.

## Methods

### Cell culture, transfection, and immunoprecipitation

HEK293T/17 verified cells purchased from ATCC (CRL-11268; STR profile, amelogenin: X) were maintained in Dulbecco’s Modified Eagle’s Medium (DMEM, ATCC) with 10 % fetal bovine serum (FBS, Gibco) at 37 °C in a humidified incubator in the presence of 5 % CO_2_. Low passage cells (less than 20 passages) were used for transfection experiments and were monitored for mycoplasma contamination using the Universal Mycoplasma Detection Kit (ATCC, 30–1012K).

Sequence encoding *gjd1a/cx34.1* fused in-frame to the V5-TurboID encoding sequence^60,61^ was constructed into a pCMV expression vector using Gibson cloning. The V5-TurboID was placed internally (following Asp278 in Cx34.1) to preserve availability of the PDZ binding domain^66^. The plasmid directing expression of mVenus-Tjp1b/ZO1b-Beta-8XHIS was generated in a previous study^83^. Low passage HEK293T/17 cells were seeded at a density of 1 × 10^6^ cells/well of a six-well dish 24 hours prior to transfection. Plasmids were co-transfected using Lipofectamine 3000 (Invitrogen) following the manufacturer’s instructions. If needed, empty pCMV plasmid was co-transfected to preserve the total concentration of transfected plasmid DNA, as denoted in the figure legend. Prior to collection 36–48 hours post-transfection, cells were incubated with culture medium supplemented with 0.5mM biotin at 37 °C in a humidified incubator in the presence of 5 % CO_2_. Cells were rinsed with 1x PBS, collected into a 1.5ml Eppendorf, and pelleted by centrifugation at 5,000 x g for 5 min at 4°C. Pellets were lysed in 0.20 ml solubilization buffer (50 mM Tris [pH7.4], 100 mM NaCl, 5 mM EDTA, 1.5 mM MgCl2, 1% Triton X-100) plus a protease inhibitor cocktail (Pierce) for 1hr with rocking at 4°C and centrifuged at 20,000 x g for 30 min at 4°C. Equal amounts of extract were immunoprecipitated with 2.0 µl rabbit anti-GFP (Abcam, ab290) overnight with rocking at 4°C. Immunocomplexes were captured with 25 µl prewashed Protein A/G agarose beads for 1 hour with rocking at 4°C, washed three times with lysis buffer, and boiled for 5 min in the presence of LDS-PAGE loading dye containing 200 mM DTT. Samples were resolved by SDS-PAGE using an 8–16% gradient gel and analyzed by Western blot using rabbit anti-GFP (Invitrogen, 11122) followed by IRDye 680LT goat anti-rabbit secondary antibody (LI-COR), IRDye 680LT (LI-COR) conjugated-rabbit anti-Cx34.1-3A4 (Fred Hutch Antibody Technology Facility, clone 3A4), and Strep-IRDye 800CW (LI-COR) and visualized with the near-infrared Odyssey system (LI-COR).

### Zebrafish care

With approval from the Institutional Animal Care and Use Committee (IACUC AUP 21-42), zebrafish (*Danio rerio*) were bred and maintained in the University of Oregon fish facility at 28 °C on a 14 hours on and 10 hours off light cycle. Animals were housed in groups according to genotype, with no more than 25 animals per tank. Development time points were assigned according to standard developmental staging^117^. All fish used for this project were maintained in the ABC background developed at the University of Oregon, and most contained the enhancer trap transgene *Et(T2KHG)^zf206^* unless otherwise noted^98^. All immunohistochemistry was performed at 5 dpf, at which stage zebrafish sex is indeterminate^72^.

### Genome engineering of *gjd1a-V5-TurboID* fish by homologous recombination

A single guide RNA (sgRNA) targeting exon 2 of *gjd1a/cx34.1* was designed using the CRISPRscan algorithm^118^ and synthesized as previously described^62^ using the T7 megascript kit (ThermoFisher). The CRISPR target sequence used to create *gjd1a-V5-TurboID* (PAM underlined) is: 5’-GCGGCCCAAGTTGGGCACGG<u>AGG</u> -3’.

A plasmid was built to contain coding sequence for the V5 affinity tag and TurboID^60,61^, flanked by homologous arms cloned from genomic DNA of ABC fish carrying the enhancer trap transgene *Et(T2KHG)^zf206^.* The 5’ homologous arm was 418bp (Chr5: 36,996,725-36,997,142) and the 3’ homologous arm was 297bp (Chr5: 36,997,143-36,997,439). Two silent mutations were placed into the CRISPR target site of the repair template (5’-GCGGCCCAAGTTGGGCACtGAtG) to prevent re-cleavage of the repaired site. A double-stranded DNA (dsDNA) repair template was generated by PCR from the plasmid template using primers modified with 5’-biotin and phosphorothioate bonds (denoted by *) for the first five nucleotides of the primer: FP: 5’ Biotin-C*A*A*A*G*ACGACCGCGAATGTCTG-3’; RP: 5’ Biotin-G*T*A*A*C*ATGAGCCCCTGCTACGTTC-3’. The 1723 bp amplicon was treated with DpnI at 37 °C for 30 min, then 80 °C for 20min, column purified and resuspended in RNase-free water.

Injection mixes were prepared in a pH 7.5 buffer solution of 300 mM KCl and 4 mM HEPES and contained a final concentration of 12 μM sgRNA, 10uM Cas9 protein (IDT), and 25nM dsDNA template. Injection mixes were incubated at 37 °C for 5 min immediately prior to injection to promote formation of the Cas9 and sgRNA complex. Parent fish containing the enhancer trap transgene *Et(T2KHG)^zf206^* were previously sequenced for those containing exact homologous arms. Embryos generated from this select parent pool were injected with 1 nL at the one-cell stage^119^. Samples of injected sibling embryos were either processed for immunofluorescence to confirm mosaic expression of V5 at M/CoLo electrical synapses or harvested for PCR analysis of genomic DNA to confirm template integration. The remaining injected embryos were raised to adulthood and outcrossed to wild-type animals. An animal carrying the homologously recombined V5-TurboID in-frame insertion was identified by PCR and immunofluorescent staining of the V5-tagged allele, and the allele was verified by Sanger sequencing *Gt(gjd1a-V5-TurboID^b1445^)*.

### Preparation of Larval Fish

Heterozygous *Gt(gjd1a-V5-TurboID^b1445^)* containing the enhancer trap transgene *Et(T2KHG)^zf206^* were in-crossed, grown to adulthood, and genotyped to identify homozygous and wildtype siblings. Homozygous *Gt(gjd1a-V5-TurboID^b1445^)* were crossed with wild type ABC fish to yield heterozygous offspring. The wild type siblings were crossed with wild type ABC fish to yield offspring for control comparisons. Crosses were set up between only one male and one female to produce a clutch, such that several siblings from each clutch could be sacrificed to verify the genotype of the entire clutch. At day 4 dpf larvae were switched to embryo medium containing 1mM biotin. At day 6 dpf larvae were euthanized with tricaine methanesulfonate (ms-222), collected, extensively rinsed in embryo medium (supplemented with ms-222) to remove exogenous biotin, and pooled together by genotype. Larvae (at least 1200 per genotype) were flash frozen in liquid nitrogen and stored at -80 °C until use. Larval preparations were collected on three separate dates.

### Preparation of Adult Fish Brains

Heterozygous *Gt(gjd1a-V5-TurboID^b1445^)* containing the enhancer trap transgene *Et(T2KHG)^zf206^* were in-crossed, grown to adulthood, and genotyped to identify heterozygous and wildtype siblings. Six males and six females of each genotype (11.5 months old siblings) were sorted into individual static tanks containing fish water supplemented with 1mM biotin. At 74 hours post-biotin, fish were euthanized and fin clipped for subsequent genotype verification. Brains were harvested, snap frozen in liquid nitrogen, and stored at -80 °C until use^120^.

### Preparation of Homogenates

Frozen larvae for each genotype were resuspended in 4mL 1X RIPA (50mM Tris pH 7.4, 150mM NaCl, 1mM EDTA, 1mM EGTA, 0.1 % SDS, 0.5 % sodium deoxycholate, 1.0 % NP-40 substitute, 1mM PMSF, and protease inhibitors [ThermoFisher A32955]) in a 5ml LoBind eppie. Samples were sonicated for 15 sec at 35% amplitude (1 sec on, 1 sec off) on ice, resting 15 sec on ice and repeated twice (total of 45 sec). Samples rotated overnight at 4 °C.

For adult brains, two brains were removed from the -80 °C freezer and pooled together as one biological sample, with a total of three biological replicates for each genotype/sex combination. The two brains were homogenized using a glass Dounce homogenizer in 750ul Homogenization buffer (50mM Tris pH 7.4, 150mM NaCl, 1mM EDTA, 1mM EGTA, 1mM PMSF, and protease inhibitors [ThermoFisher A32955]). The homogenate was transferred to a 2mL LoBind eppie and an equal volume of 2X RIPA buffer (50mM Tris pH 7.4, 150mM NaCl, 1mM EDTA, 1mM EGTA, 0.2% SDS, 1.0% sodium deoxycholate, 2.0% NP-40 substitute, 1mM PMSF, and protease inhibitors [ThermoFisher A32955]) was added (final concentration of detergents: 0.1% SDS, 0.5% sodium deoxycholate, 1.0% NP-40). Samples were sonicated for 15 sec at 35% amplitude (1 sec on, 1 sec off) on ice. Samples rotated overnight at 4 °C.

The following day samples were centrifuged 14,000 x g, 20min, 4°C. The supernatant was desalted over a PD-10 column (Cytiva) pre-equilibrated with 1X RIPA buffer (50mM Tris pH 7.4, 150mM NaCl, 1mM EDTA, 1mM EGTA, 0.1 % SDS, 0.5 % sodium deoxycholate, 1.0 % NP-40 substitute) following the manufacturer’s protocol. The protein concentration of the eluent was determined using the Pierce BCA Protein Assay Kit. Equal amounts of total protein were aliquoted into a 5mL LoBind eppie and prewashed Streptavidin magnetic beads (Pierce Streptavidin Magnetic Beads, Cat# 88817) were added. For larvae, three separate dates of collection represent the three biological replicates for each genotype. Collection date #1 contained one replicate for each genotype (15 mg total protein/300ul prewashed Streptavidin magnetic beads). Collection dates #2 and #3 contained two technical replicates for each genotype (7mg total protein/150ul prewashed Streptavidin magnetic beads). For adults, one date of collection represents the three biological replicates for each genotype for each sex (1mg total protein/150ul prewashed Streptavidin magnetic beads). Samples were rocked overnight at 4 °C.

The following day, magnetic streptavidin beads were washed using a magnetic stand once with wash buffer 1 (50mM Tris pH 7.4, 2% SDS), twice with wash buffer 2 (50mM Tris pH 7.4, 150mM NaCl, 1mM EDTA, 0.1% SDS, 0.5% sodium deoxycholate, 1.0% NP-40 substitute), once with wash buffer 3 (50mM Tris pH 7.4, 150mM NaCl, 1mM EDTA, 1.-% NP-40), and three times with wash buffer 4 (50mM ammonium bicarbonate in mass spectrometry grade water). Samples were immediately shipped on wet ice overnight to the OHSU Proteomics Shared Resource mass spectrometry facility for analysis.

### Mass Spectrometry

Magnetic streptavidin bead samples were washed twice with 100ul of 100mM ammonium bicarbonate and resuspended in a final volume of 100ul. Trypsin (12.5ul of 80ng/μl trypsin in 50mM TEAB (1ug) was added to each sample, mixed gently and incubated at 37C overnight with shaking on a Thermomixer at 800rpm. The following day samples were placed on a magnetic stand for 2min, the supernatant removed, filtered through a 0.22um Millipore filter and dried. The dried filtered sample was dissolved in 20ul of 5% Formic acid and injected into QExactive HF and run with a 90min LC/MS method. Steps were performed at the Oregon Health Sciences University (OHSU) Proteomic Shared Resource.

### Mass Spectrometry Data Analysis

The binary instrument files were processed with the PAW pipeline (https://github.com/pwilmart/PAW_pipeline)^121^. Binary files were converted to text files using MSConvert^122^. Python scripts extracted fragment ion spectra in MS2 format^123^. The Comet search engine (version 2016.03)^124^ was used: 1.25 Da monoisotopic peptide mass tolerance, 0.02 Da monoisotopic fragment ion tolerance, fully tryptic cleavage with up to three missed cleavages, variable oxidation of methionine residues, and static alkylation of cysteines. Searches used UniProt proteome UP000000437 (zebrafish, taxon ID 7955) canonical FASTA sequences (20,603 proteins). Six additional protein sequences and common contaminants (174 sequences excluding any albumins) were added, and sequence-reversed entries were concatenated for a final protein FASTA file of 52,824 sequences (https://github.com/pwilmart/fasta_utilities).

Top-scoring peptide spectrum matches (PSMs) were filtered to a 3% false discovery rate (FDR) using interactive delta-mass and conditional Peptide-prophet-like linear discriminant function scores^125^. Incorrect delta-mass and score histogram distributions were estimated using the target/decoy method^126^. The filtered PSMs were assembled into protein lists using basic and extended parsimony principles and required two distinct peptides per protein. The final list of identified proteins, protein groups, and protein families were used to define unique and shared peptides for quantitative use. Total (summed) corrected spectral counts were computed for each protein where shared peptide counts were fractionally split between the proteins they mapped to based on relative unique count totals. The overall protein FDR was 1.1%.

The protein corrected spectral count values for each biological sample in each biological condition were compared for differential protein expression using the Bioconductor package edgeR^127^ exact test within Jupyter notebooks. For developmental samples or adult brain samples, a minimum average spectral count of 1.5 was required for quantification. A count value of one was added to all spectral count totals for each protein to remove any zero count values. Result tables contained typical proteomics summaries, spectral count totals, and statistical testing results.

To aid interpretation of results, the zebrafish identified protein sequences were matched against the human canonical reference proteome use BLAST and Python scripts (https://github.com/pwilmart/PAW_BLAST and https://github.com/pwilmart/annotations). Additional UniProt annotation information was added for the human ortholog proteins (GO terms and Reactome pathways). STRING and g:Profiler were used for Gene Ontology (GO) analysis^76,77^. Resources including SFARI^78^, OMIM^79^, and Genecards^80^ that catalog human-disease-associated genes were used to for gene annotation.

### scRNA-seq analysis

The single-cell RNA sequencing (scRNAseq) data used in this study originates from the Farnsworth et al. 2020 dataset (5 dpf)^73^ and the Posner et al. 2024 dataset (6 dpf)^128^. Both datasets were processed using the pipeline described in Lukowicz-Bedford et al. 2022^67^. Briefly, pooled larval samples were collected in two replicates at each time point. Standard protocols for cell dissociation from whole larvae were followed^73^. The dissociated cells were processed on the 10X Chromium platform using v2 chemistry for the 5 dpf dataset and v3 chemistry for the 6 dpf dataset, targeting approximately 10,000 cells per run. Aligned reads were mapped to the zebrafish genome (GRCz11) using the 10X Cellranger pipeline (version 3.1), with an updated GTF file that captures Connexin-encoding gene expression^67^. Cells were processed in Seurat (v5) for R (v4.3.2) with standard quality control, normalization, integration, and analysis procedures^129^. Analysis code and data sets can be found at www.adammillerlab.com/resources.

Germ layer clusters were identified using established tissue markers from Farnsworth et al. 2022. Mesoderm clusters included putative skeletal muscle, blood, and kidney cells (markers: *myod1, smyhc1, myhz2, tnnc1b, gata1a, hbae1.1, cahz, slc4a1a, slc5a9, prr15lb, aqp8b, pdzk1ip1*). Endoderm clusters included putative liver cells (*foxa3, fabp10a, prox1a, cp*). Non-neural ectoderm clusters included putative skin and cranial neural crest cells (*trpv6, grhl1, gjb8, foxi3a, sox10, dlx2a, foxd3, crestin*). Neural ectoderm clusters were identified based on neural markers (*elavl3, elavl4, snap25a*).

### Cas9-mediated genome engineering of *V5-tagged* mosaics

The CRISPR target sequences used to insert V5 coding sequence for each gene (PAM underlined) are noted in Table S3. The V5-tagged single-stranded donor oligos (ssODN) were designed to repair into endogenous loci, as listed to Table S3. The ssODN contained ∼24-36 bp homology arms, tandem V5 sequences, and a 5x-glycine linker between V5 tags and between the V5 tags and endogenous gene sequence. If the inserted sequence did not disrupt the endogenous sgRNA recognition site, silent mutations were designed in the CRISPR/PAM sites of the ssODN to prevent further double stranded breaks after repair.

Injection mixes were prepared in a pH 7.5 buffer solution of 300 mM KCl and 4 mM HEPES and contained a final concentration of 2uM ssODN, 100-1000pg sgRNA, and 8 μM Cas9 protein (IDT) and were incubated at 37°C for 5 min prior to injection to promote formation of the Cas9 and sgRNA complex. Embryos containing the enhancer trap transgene *Et(T2KHG)^zf206^* were injected with 1 nL at the one-cell stage^95^. At 3-4 dpf, injected embryos were screened by high resolution melting (HRM) to confirm active CRISPRing using gene-specific HRM primers as denoted in Table S3. At 5 dpf, injected larvae were screened by immunohistochemistry for mosaic expression of V5-tagged proteins.

### Immunohistochemistry and confocal imaging

Anesthetized, 5 dpf larvae were fixed for 3 hours in 2% trichloroacetic acid in PBS^130^. Fixed tissue was washed in PBS containing 0.5% Triton X-100, followed by standard blocking and antibody incubations. Primary antibody mixes included combinations of the following: rabbit anti-Cx34.1 3A4 (Fred Hutch Antibody Technology Facility, clone 3A4, 1:200), mouse IgG1 anti-NF(RM044, Invitrogen, 13-0500, 1:200), mouse IgG2a anti-V5 (Invitrogen, R960-25, 1:500), mouse IgG2b anti-acetylated tubulin (Millipore Sigma, T7451-25UL, 1:1000) and chicken anti-GFP (Abcam, ab13970, 1:500). All secondary antibodies were raised in goat (Invitrogen, conjugated with Alexa-405-Plus, –488, −555, −633, or −647 fluorophores, 1:500). Tissue was cleared stepwise in a 25%, 50%, 75% glycerol series, dissected, and mounted in ProLong Gold antifade reagent (ThermoFisher). Using a Leica SP8 Confocal microscope, images were acquired using a 405-diode laser and a white light laser set to 499, 553, 598, and 631 nm, depending on the fluorescent dye imaged. Laser line data was collected sequentially using custom detection filters based on the dye. Images of the Club Endings (CEs) were collected using a 63x, 1.40 numerical aperture (NA), oil immersion lens, images of M/CoLo synapses were collected using a 40x, 1.20 NA, water immersion lens, and tile scans were collected with either objective protocol, as noted in figure legend. The optimal optical section thickness was calculated by the Leica software based on the pinhole, emission wavelengths, and NA of the lens. For whole brain images, tile scans were stitched together using the LAS X software (Leica). Within each experiment where fluorescence intensity was to be quantified, all animals were stained together using the same antibody mix, processed at the same time, and all confocal settings (laser power, scan speed, gain, offset, objective, and zoom) were identical. The M/CoLo synapses analyzed began at approximately somite 10 within the spinal cord. Similar numbers of neighboring, caudal somites were imaged and analyzed in all animals. Multiple animals per genotype were analyzed to account for biological variation, and fluorescence intensity values for each region of each animal were an average across multiple synapses to account for technical variation.

### Analysis of confocal imaging

For fluorescence intensity quantitation, FiJi software was used to process and analyze confocal images^131^. To quantify staining at CE synapses, confocal z-stacks of the Mauthner soma and lateral dendrite (centered around the lateral dendritic bifurcation) were cropped to 200 x 200 pixels. Based on GFP staining, a FIJIscript cleared the region outside of the Mauthner cell, and for each channel a standard threshold was set to remove background staining. The image was then transformed into a max intensity projection, synapse thresholds were set to WT levels, and the integrated density of each channel within the club ending synapses was measured. To quantify staining at M/CoLo synapses, a standard region of interest (ROI) surrounding each M/CoLo site of contact was drawn around the synapse, and the mean fluorescence intensity was measured. Values were normalized to the animals with the highest value in the group (denoted as a grey bar in the figure), and n represents the number of fish used. Figure images were created using FiJi and Illustrator (Adobe).

### Statistical analysis of confocal imaging

Standard deviation, standard error of the mean, and the final statistical analysis was performed using Prism (GraphPad) software. Dunnett’s post-hoc multiple comparisons test followed all ANOVA. Statistical tests performed and p values are noted in the figure legends.

## Figure Legends

**Figure S1.**
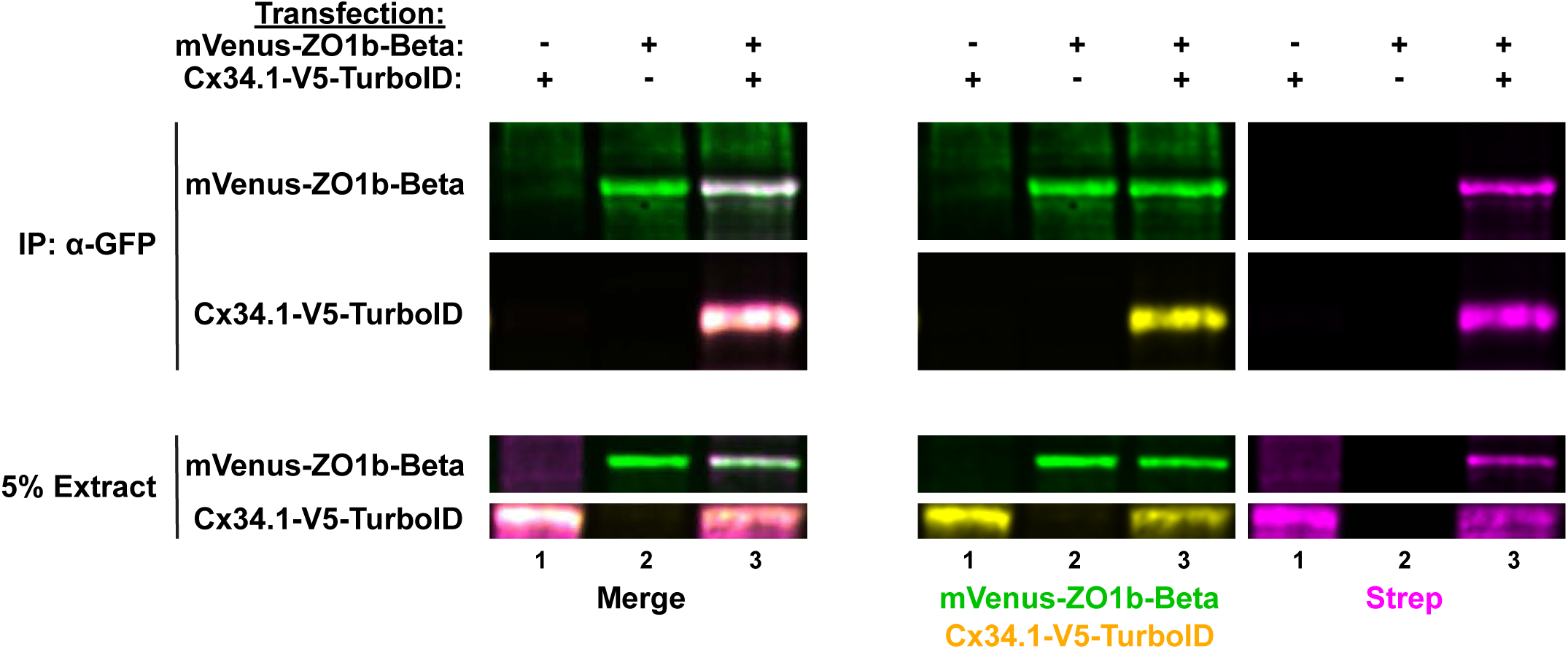
Connexin-V5-TurboID biotinylates expected proteins in cell culture. HEK293T/17 cells were transfected with plasmids to express Cx34.1-V5-TurboID or empty vector (-) together with mVenus-ZO1b (+) or empty vector (-). Prior to harvest, transfected cells were treated for 10 min with fresh cell culture medium supplemented with 1mM biotin. Lysates were immunoprecipitated with anti-GFP antibody and analyzed by immunoblot for the presence of mVenus-ZO1b using anti-GFP antibody (green), Cx34.1 protein using Cx34.1-specific antibody (yellow), or biotin using Strep-conjugated fluorophore (magenta) as shown in the channel separated images (middle and right panels). Biotinylated transfected proteins appear white in the merged images (left panels). Total extracts (bottom panels, 5% input) were blotted for transfected proteins to demonstrate uniform antibody recognition and equivalent expression of proteins. Results are representative of three independent experiments.

**Figure S2.**
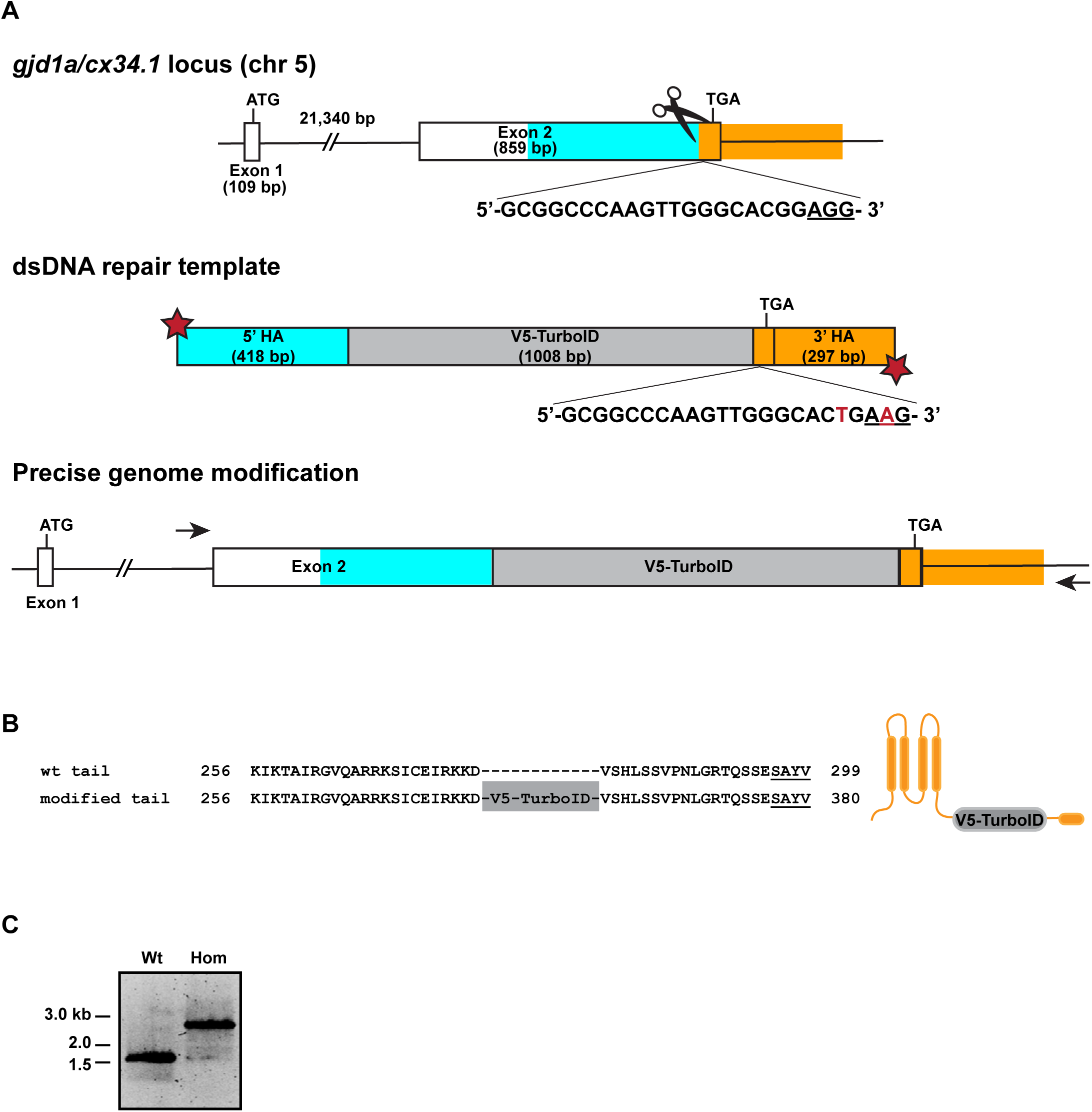
Generation scheme and verification of *gjd1a/cx34.1^V5-TurboID^* fish. **(A)** Schematic diagram of the *gjd1a/cx34.1* gene locus on chromosome 5:36,974,931-36,997238 (GRCz11 Ensembl) showing the exon structure located on the forward strand. Horizontal black bar represents the DNA strand, and boxes represent individual exons as labeled. The CRISPR location is indicated by cartoon scissors in Exon 2. The native CRISPR sequence is listed (PAM is underlined) and the silent mutations incorporated into the repair template are indicated in red font. Areas used as homologous arms (HA) in the dsDNA repair template are represented by cyan boxes (5’ HA, 418bp) and orange boxes (3’ HA, 297bp). The inserted V5-TurboID cassette (1008bp) is represented by a grey box. The red stars indicate the dsDNA template is modified by 5’ biotin plus phosphorothioate bonds between the first five nucleotides. Lengths of DNA regions are indicated. Arrows indicate the location of the flanking primers used to genotype the fish (as shown in panel C). **(B)** Amino acid sequence of the Cx34.1 wildtype C-terminal tail compared to the predicted amino acid sequence of the Cx34.1-V5-TurboID protein produced in *gjd1a/cx34.1^V5-TurboID^* fish. PDZ binding motif (PBM) is underlined. To the right, a cartoon of the Cx34.1-V5-TurboID monomer illustrates the in-frame insertion of the V5-TurboID cassette after the four trans-membrane domains (vertical orange bars) and before the C-terminal PBM (horizontal orange bar). **(C)** Genotyping of wildtype (Wt) versus *gjd1a/cx34.1^V5-TurboID^* homozygous (Hom) fish by PCR using flanking primers indicated in (A). PCR products were resolved by agarose gel and detected by SYBR Safe stain. Bands were gel purified and isolated DNA was sequenced to verify precise genome modification.

**Figure S3.**
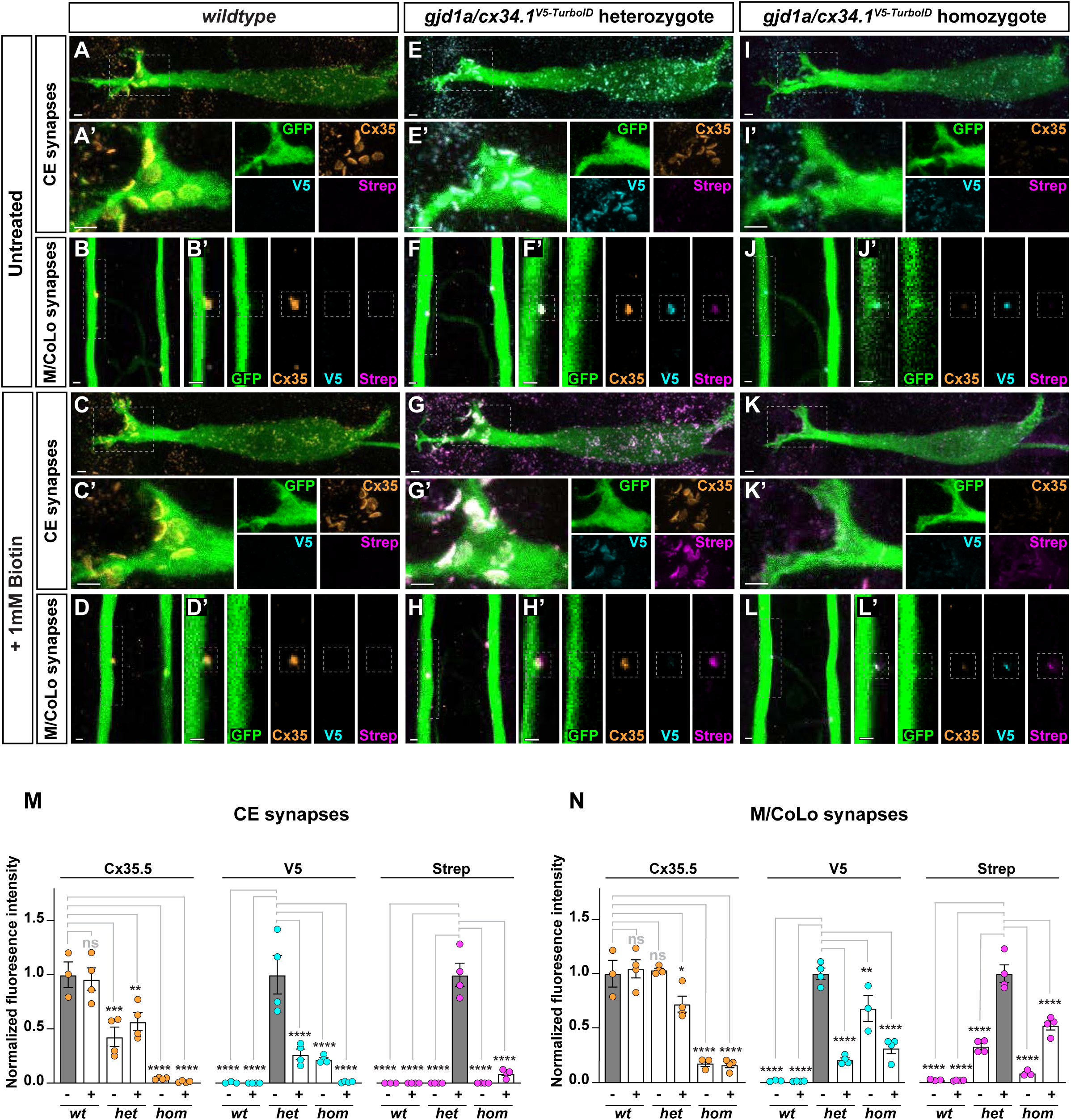
Characterization of electrical synapse biotinylation in *gjd1a/cx34.1^V5-TurboID^* fish. **(A-L)** Confocal images of M-circuit neurons and stereotypical electrical synaptic contacts in 6-day-post-fertilization, *Et(T2KHG)^zf206^* zebrafish larvae from *wildtype* (A-D), *gjd1a/cx34.1^V5-TurboID^* heterozygous (E-H) and *gjd1a/cx34.1^V5-TurboID^* homozygous (I-L) siblings either untreated (A-J) or treated for 72 hours with 1mM biotin (C-L). Animals are stained with anti-GFP (green), anti-Cx35.5 (orange), anti-V5 (cyan), and Strep-conjugated fluorophore (magenta). Scale bar = 2 µm in all images. Anterior up. Boxed regions denote stereotyped location of electrical synapses and regions are enlarged in neighboring panels. Images of the M-cell body and lateral dendrite in the hindbrain (A, C, E, G, I, K) are maximum intensity projections of ∼20-27 µm. In A’, C’, E’, G’, I’, and K’, images are maximum-intensity projections of ∼12-15 µm and neighboring panels show the individual channels. Images of the sites of contact of M/CoLo processes in the spinal cord (B, D, F, H, J, L) are maximum-intensity projections of ∼6-7 µm. In B’, D’, F’, H’, J’ and K’, images are from a single 0.42 µm Z-plane and the white dashed square denotes the location of the M/CoLo site of contact. Neighboring panels show individual channels. **(M)** Quantification of Cx35.5 (orange), V5 (cyan), and Strep (magenta) fluorescence intensities at CE synapses for the noted genotypes. The height of the bar represents the mean of the sampled data normalized to the genotype indicated by the grey bar. Circles represent the normalized value of each individual animal. Mean is shown ± SEM. For untreated CE synapses, *wt* n=3; *gjd1a/cx34.1^V5-TurboID^* heterozygotes n=4, *gjd1a/cx34.1^V5-TurboID^* homozygotes n=4. For biotin treated CE synapses, *wt* n=4; *gjd1a/cx34.1^V5-TurboID^* heterozygotes n=4, *gjd1a/cx34.1^V5-TurboID^* homozygotes n=4. **** indicates p<0.0001, *** indicates p<0.0005, ** indicates p<0.0065, and ns is not significant by ANOVA with Dunnett’s test. **(N)** Quantification of Cx35.5 (orange), V5 (cyan), and Strep (magenta) fluorescence intensities at M/CoLo synapses for the noted genotypes. The height of the bar represents the mean of the sampled data normalized to the genotype indicated by the grey bar. Circles represent the normalized value of each individual animal. Mean is shown ± SEM. For untreated M/CoLo synapses, *wt* n=3; *gjd1a/cx34.1^V5-TurboID^* heterozygotes n=4, *gjd1a/cx34.1^V5-TurboID^* homozygotes n=3. For biotin treated M/CoLo synapses, *wt* n=4; *gjd1a/cx34.1^V5-TurboID^* heterozygotes n=4, *gjd1a/cx34.1^V5-TurboID^* homozygotes n=4. **** indicates p<0.0001, ** indicates p<0.0025, * indicates p<0.0399, and ns is not significant by ANOVA with Dunnett’s test.

**Figure S4.**
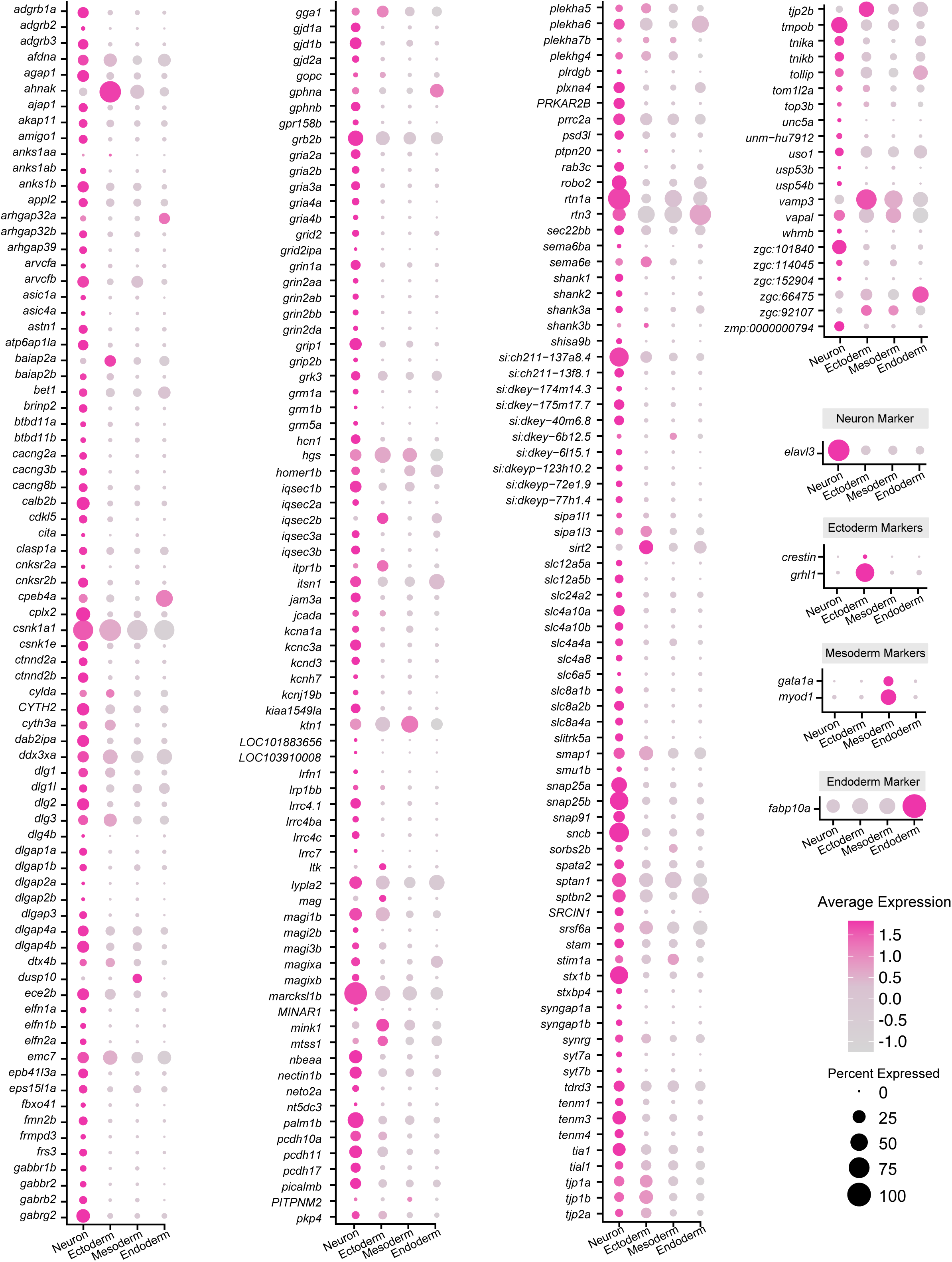
scRNA sequencing analysis of all proteome candidates demonstrates high neural expression. Neural Connexin-associated proteome gene expression across germ layers in a whole-embryo single-cell RNA sequencing dataset (5 and 6 dpf). All genes corresponding to Connexin-associated proteins are listed alphabetically along the y-axis. Clusters are organized by germ layer along the x-axis: neurons, ectoderm (non-neural ectoderm, e.g. skin and cranial neural crest), mesoderm (e.g., skeletal muscle and blood), and endoderm (e.g. liver). Marker genes for specific germ layers are highlighted: *elavl3* (neurons), *crestin* (cranial neural crest, ectoderm), *grhl1* (skin, ectoderm), *gata1a* (blood, mesoderm), *myod1* (skeletal muscle, mesoderm), and *fabp10a* (liver, endoderm). Dot size represents the percentage of cells in the category expressing each gene, while color indicates relative expression levels.

**Figure S5.**
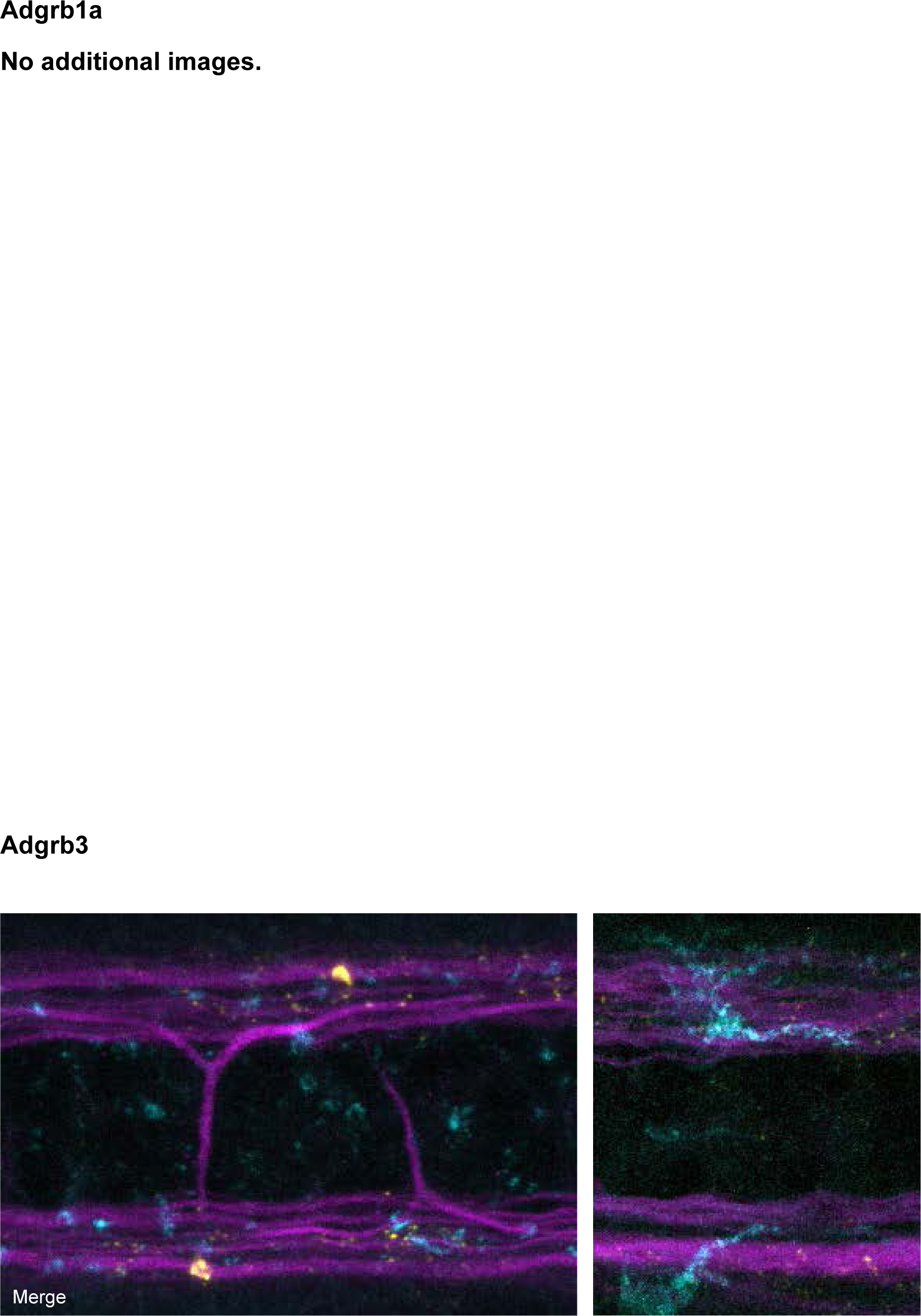
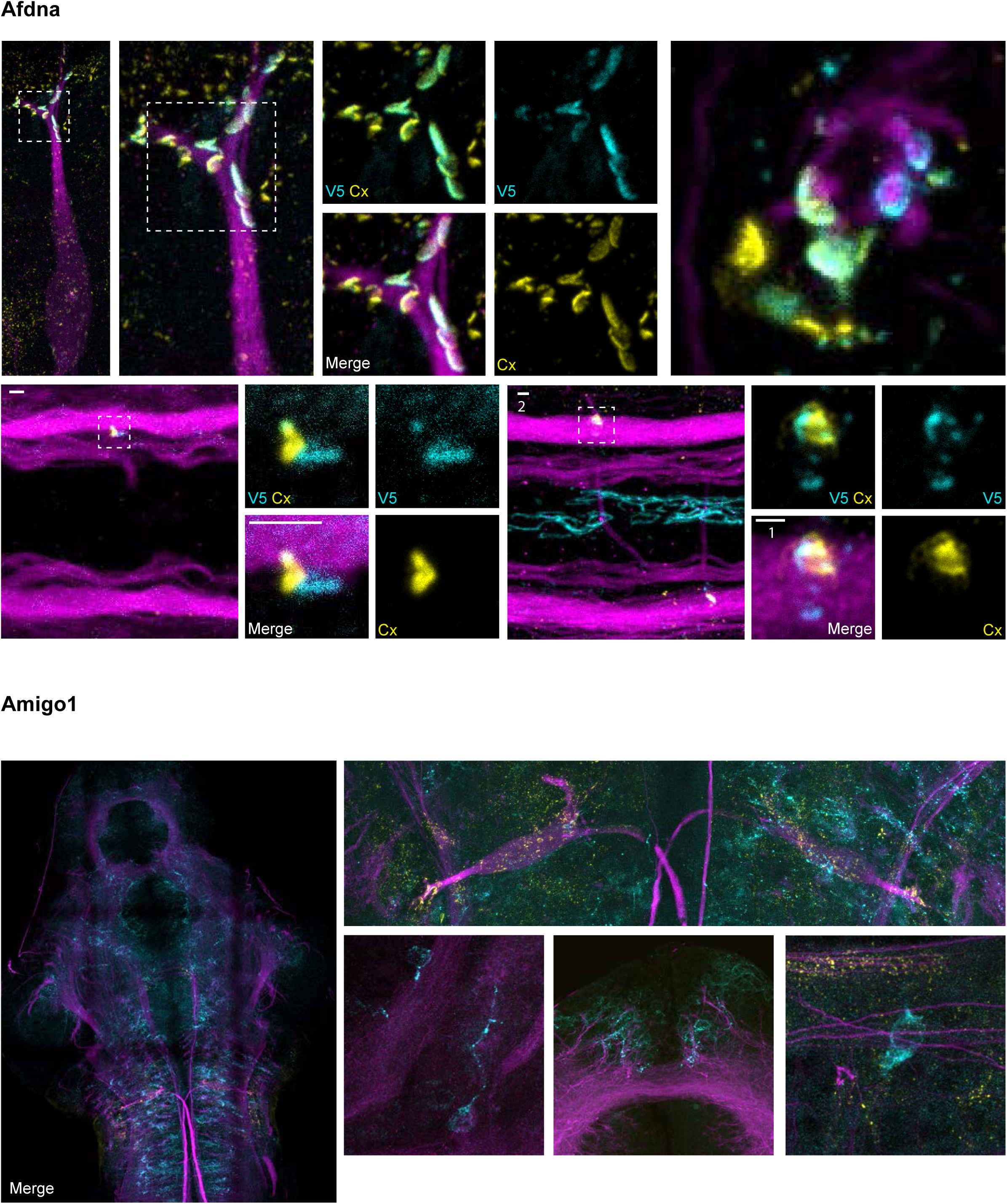
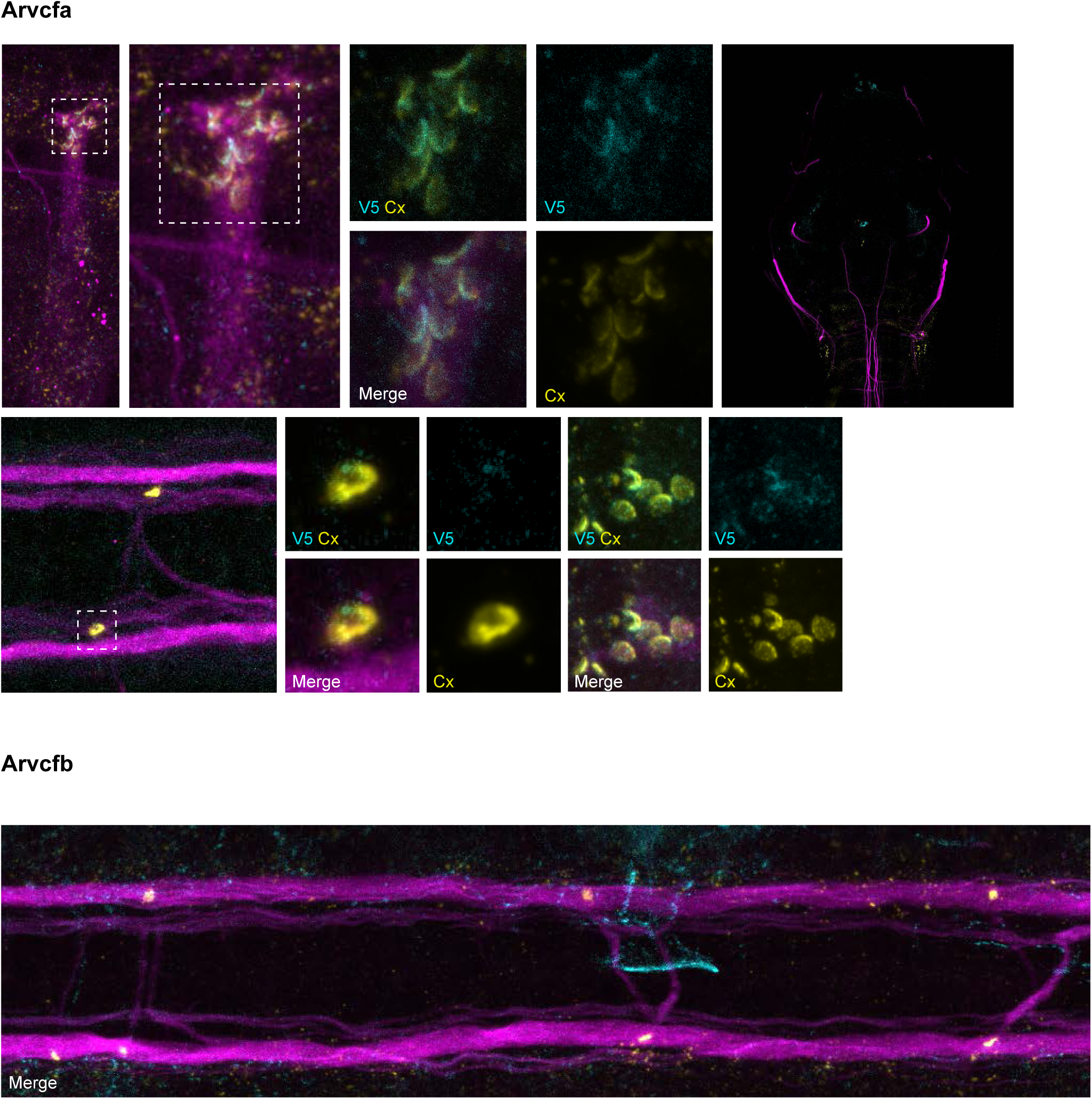
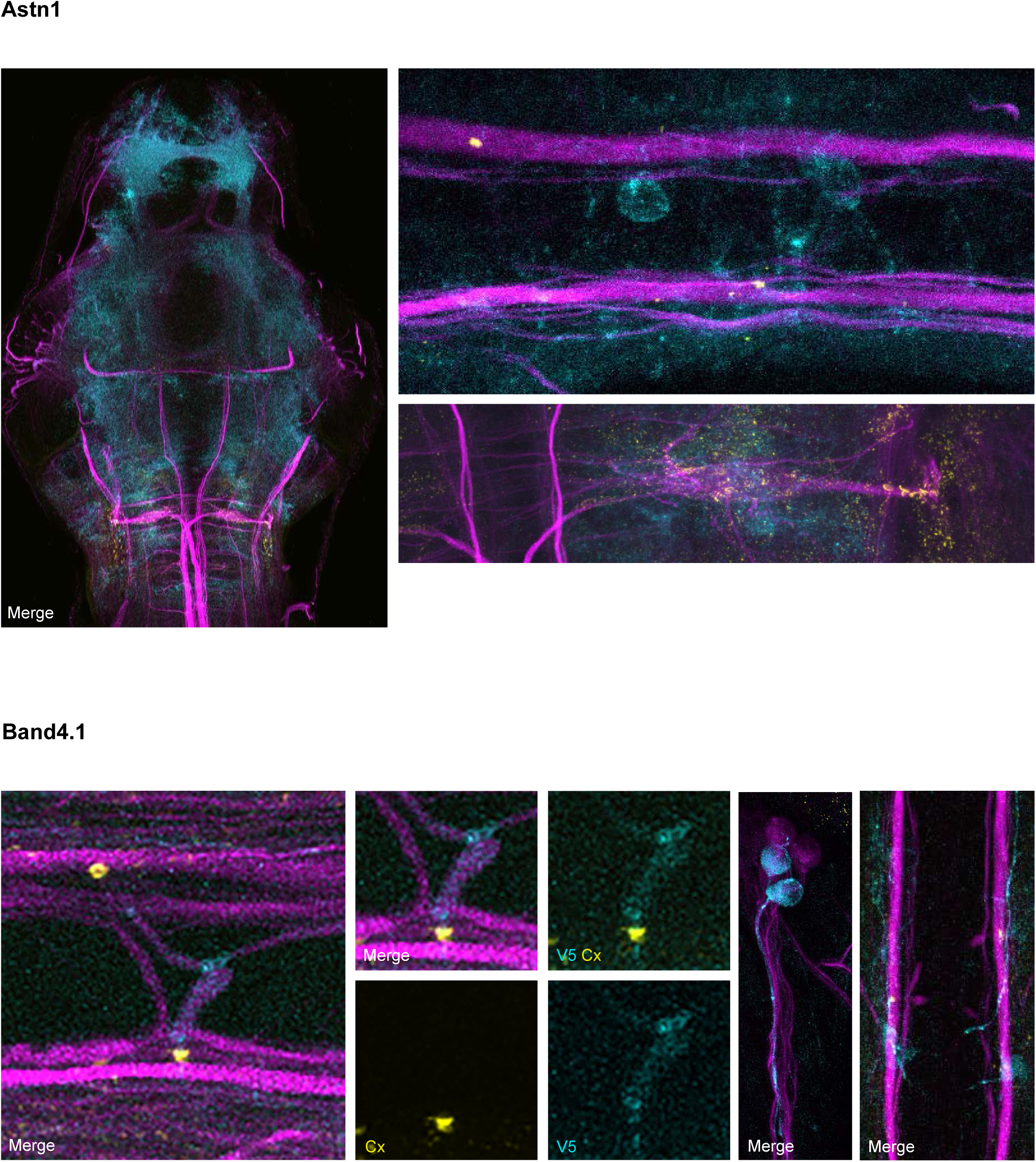
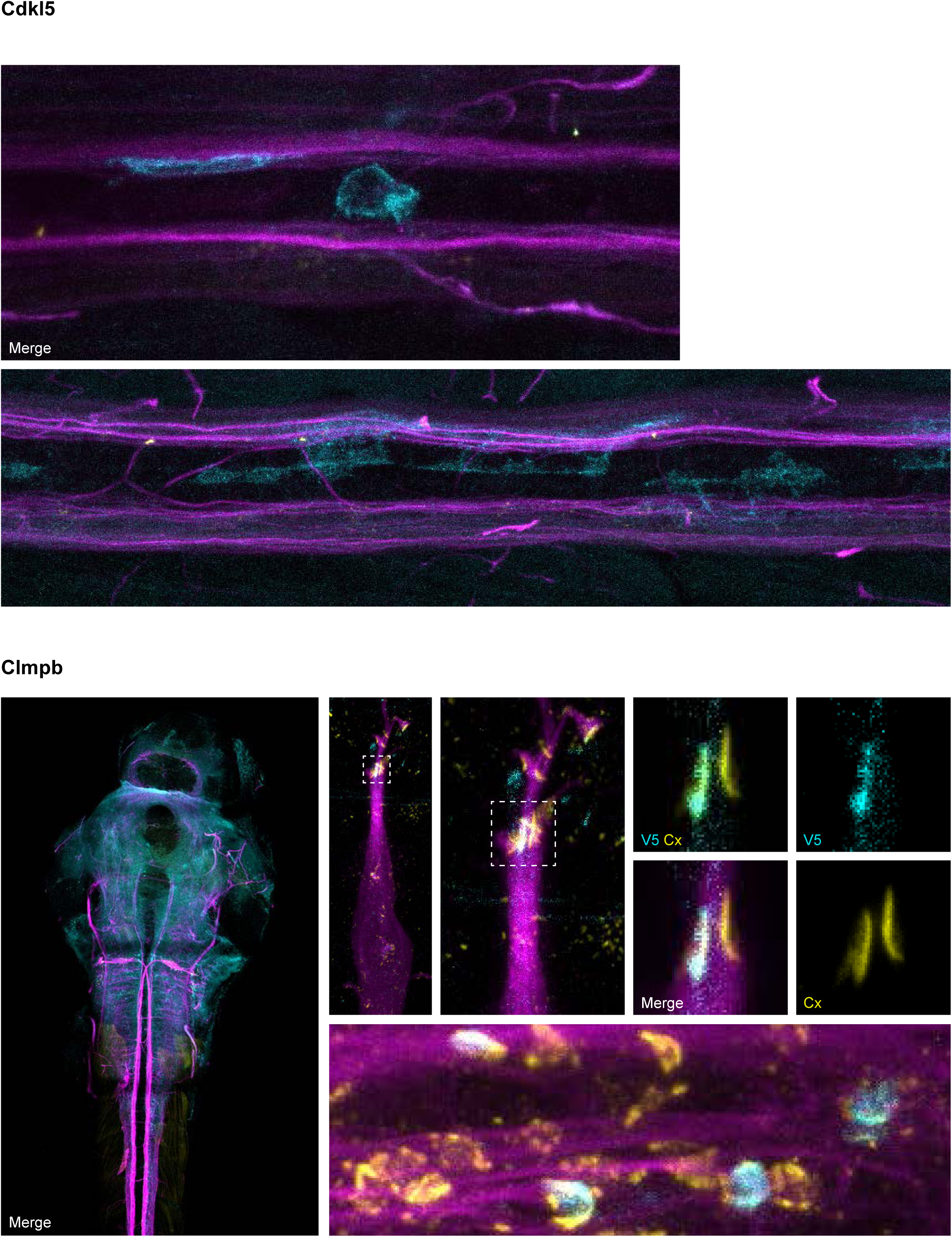
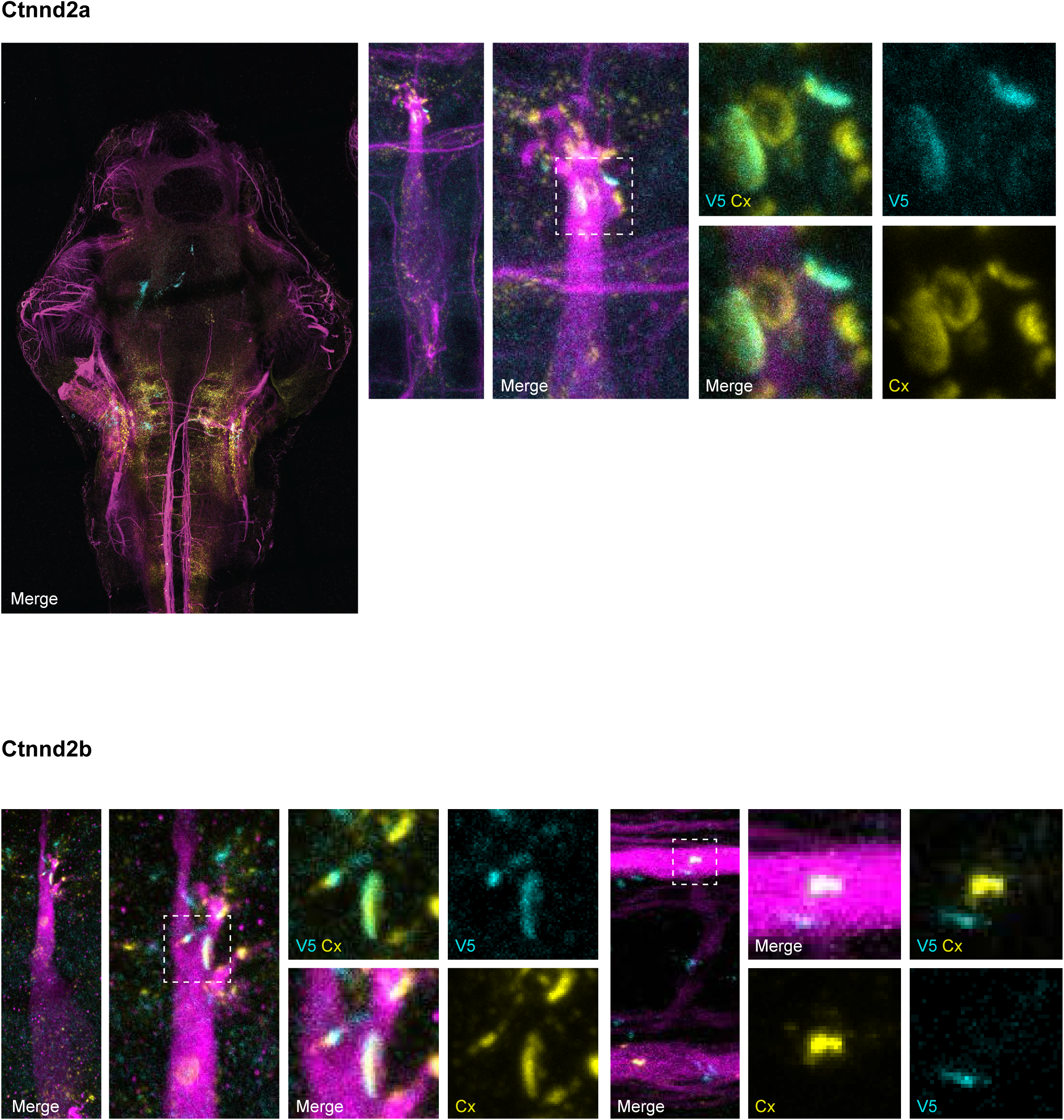
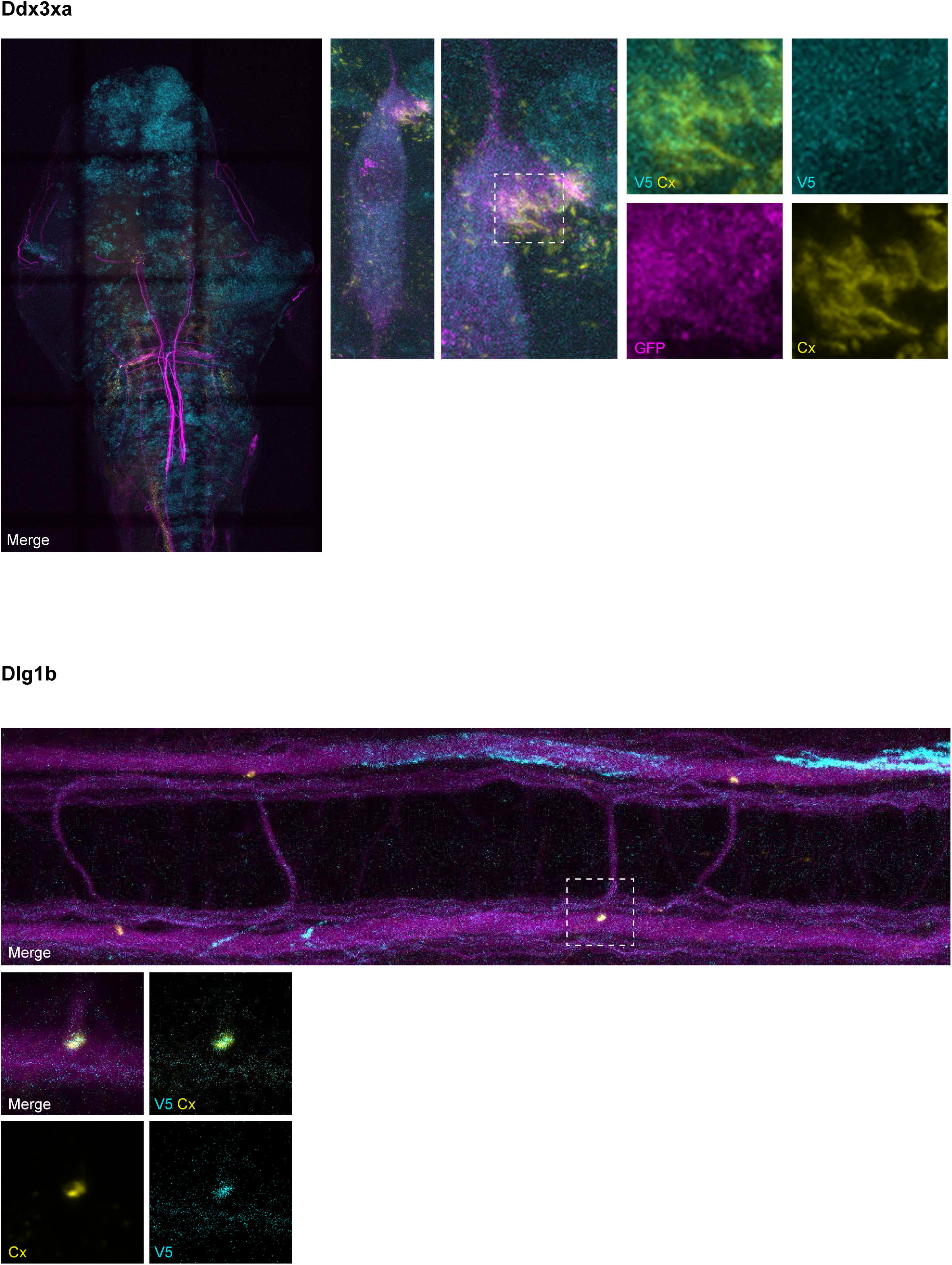
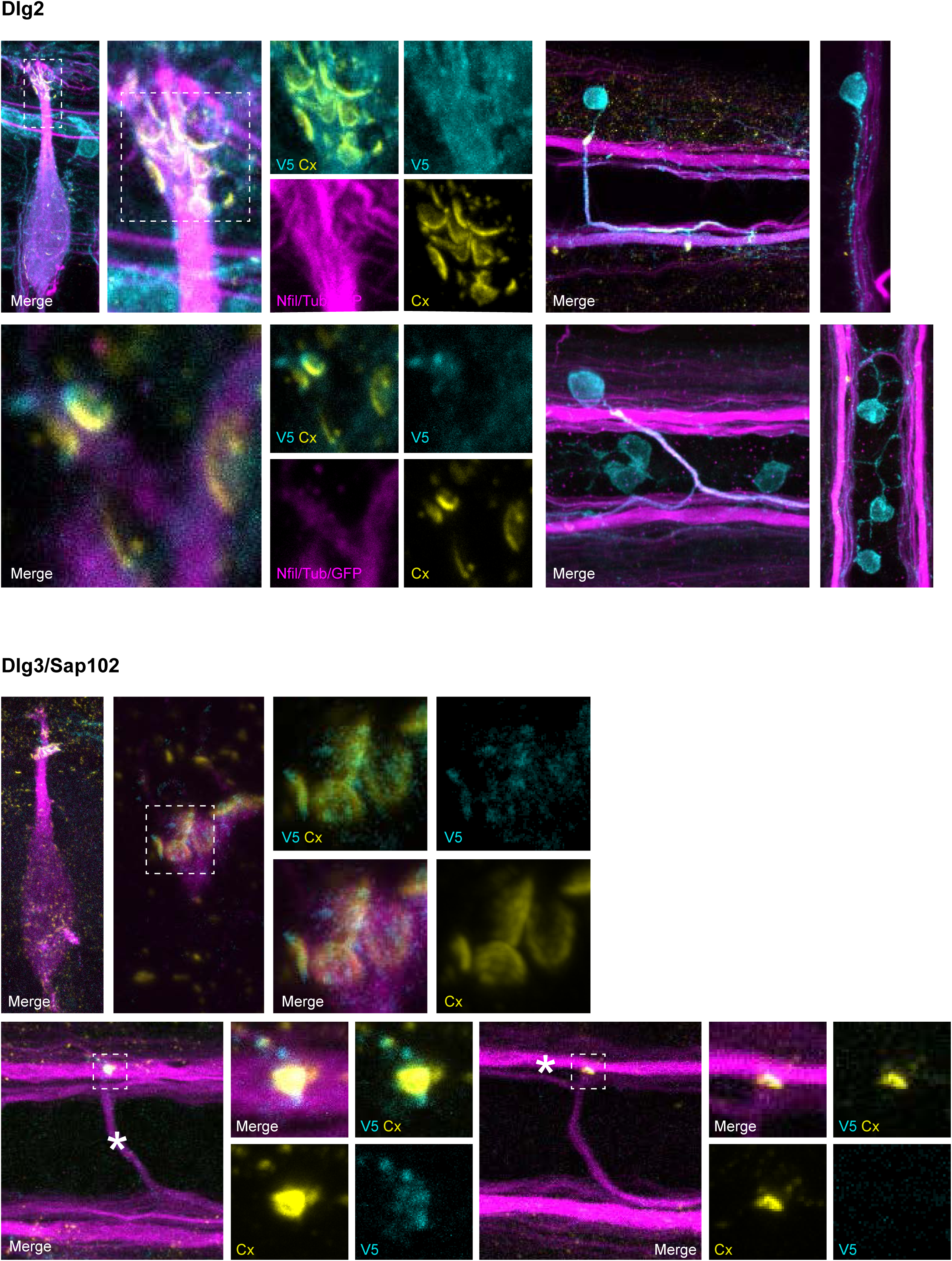
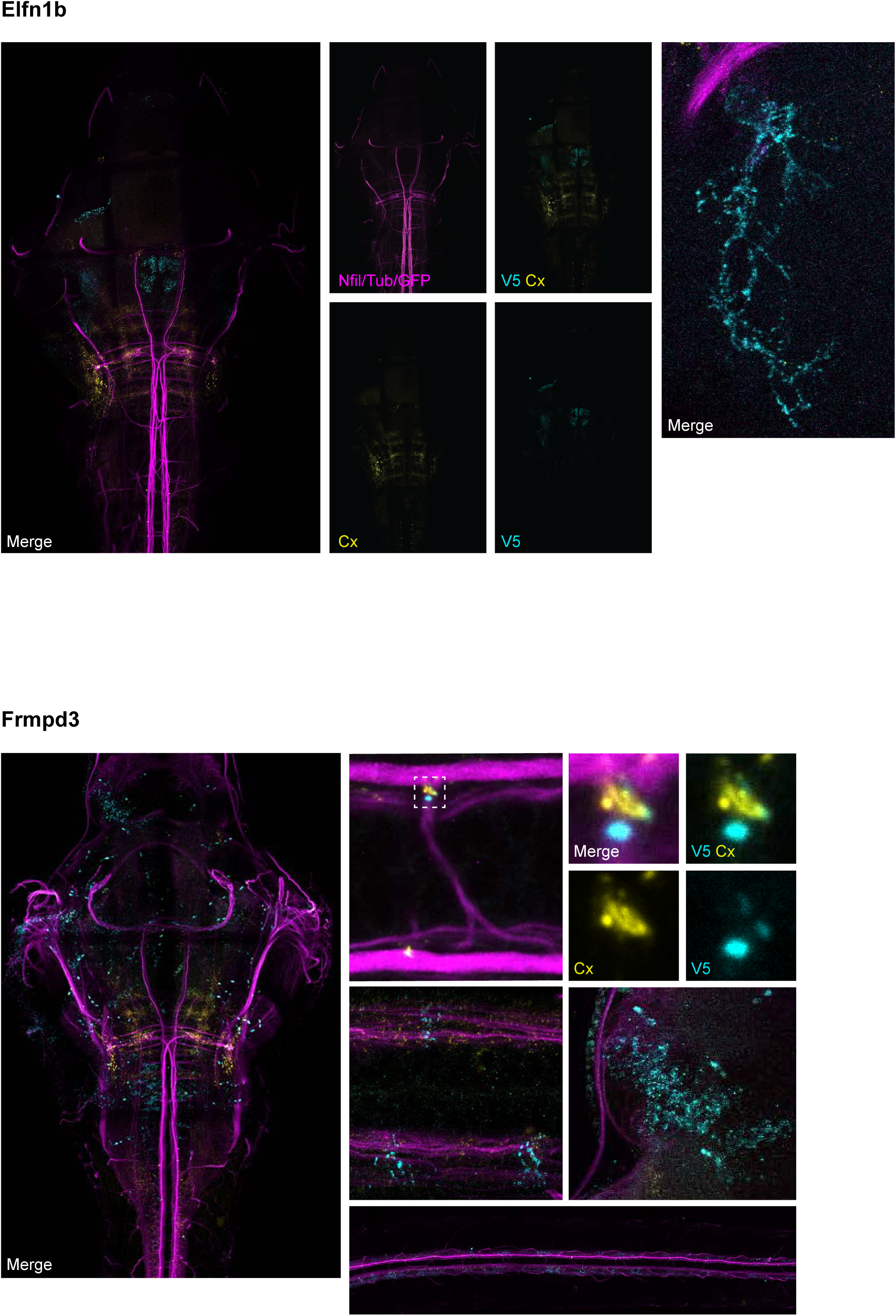
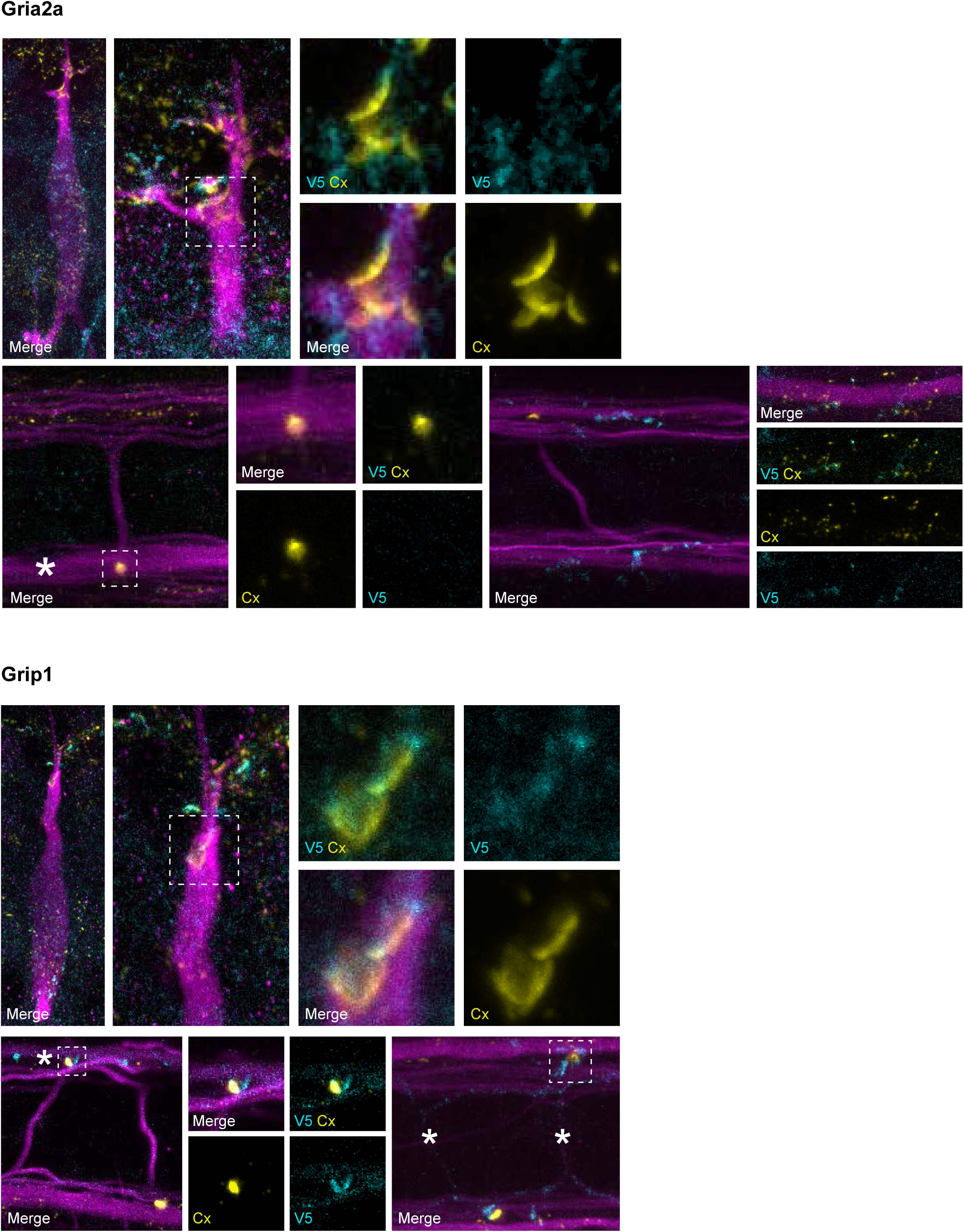
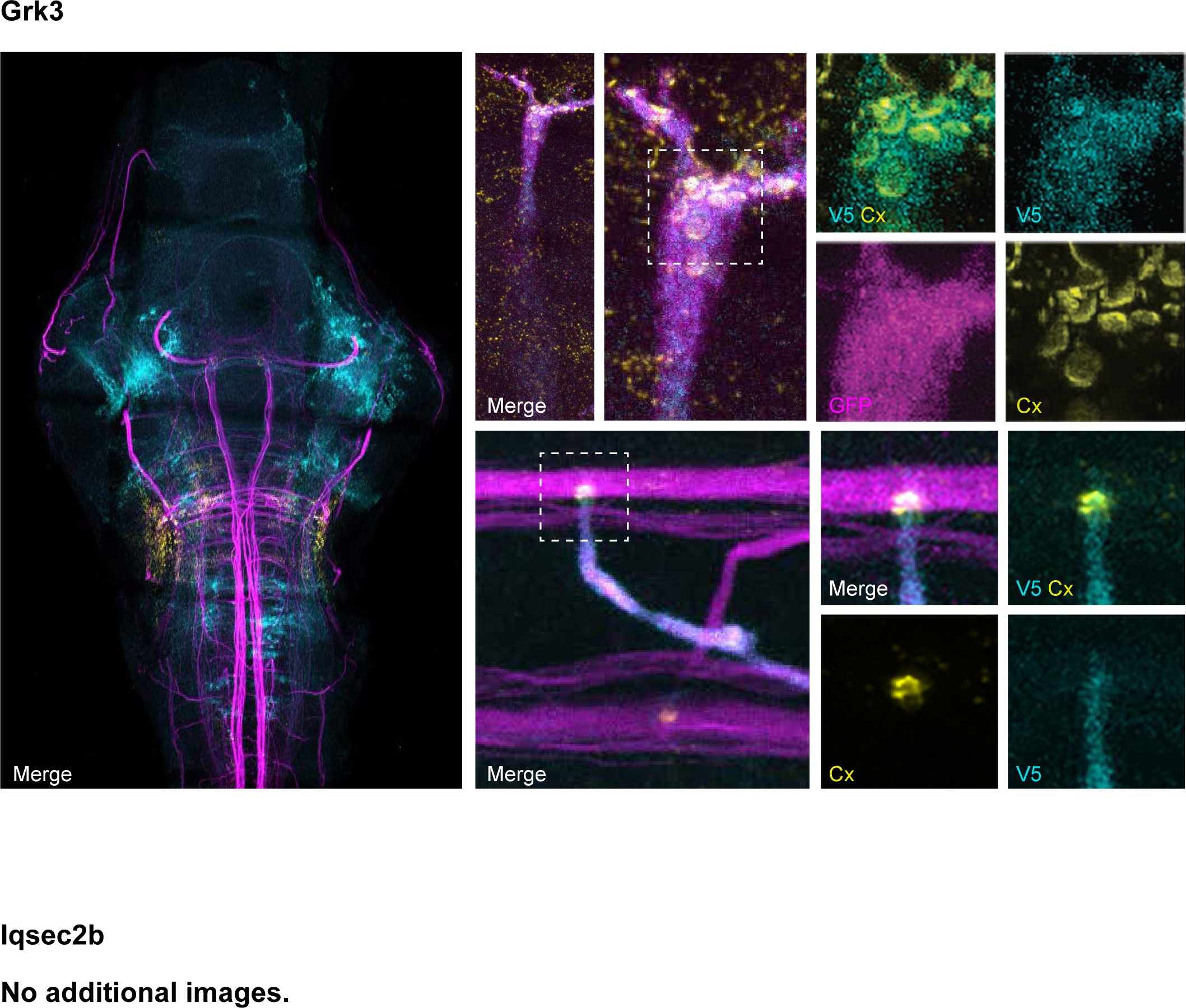
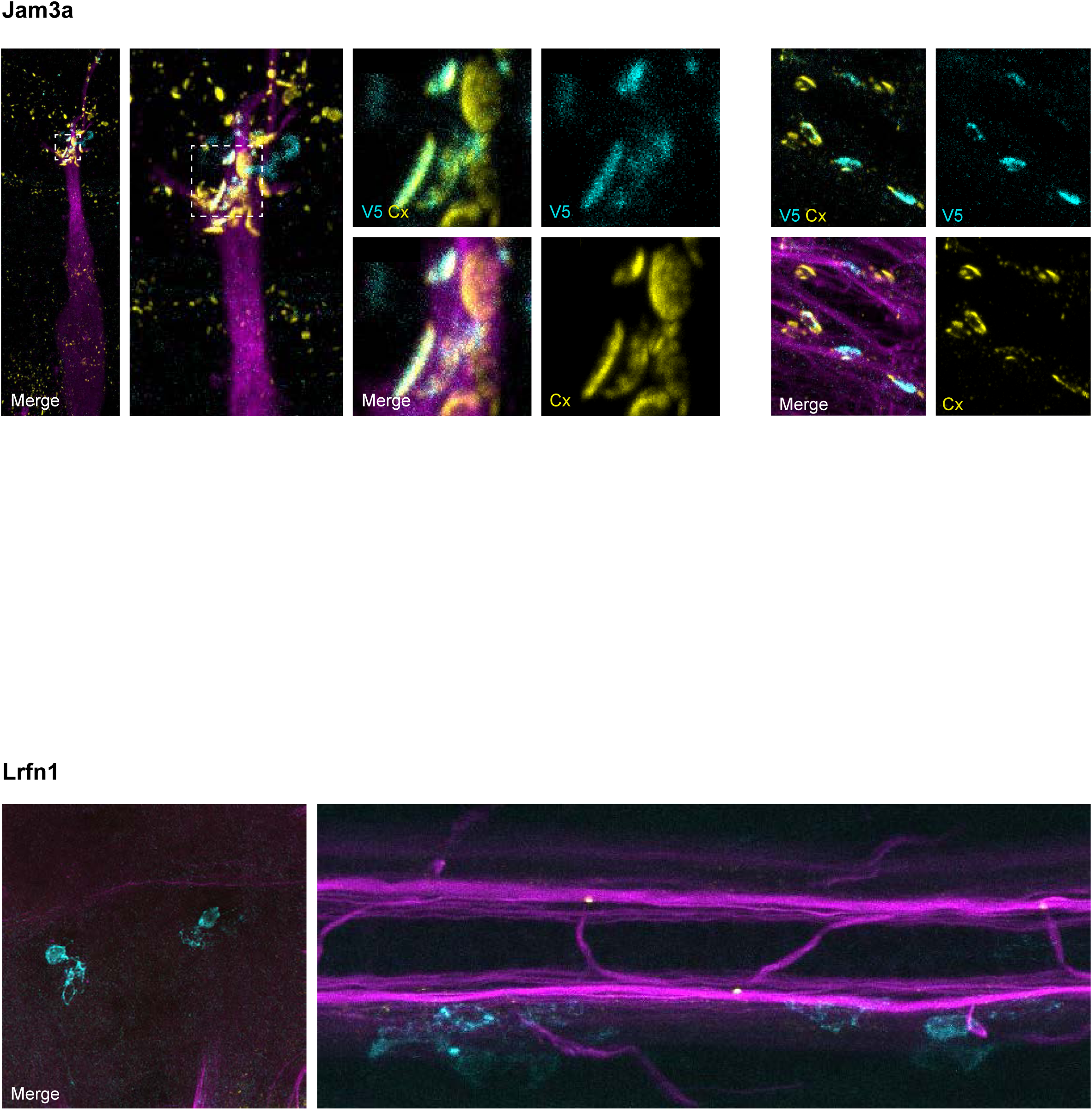
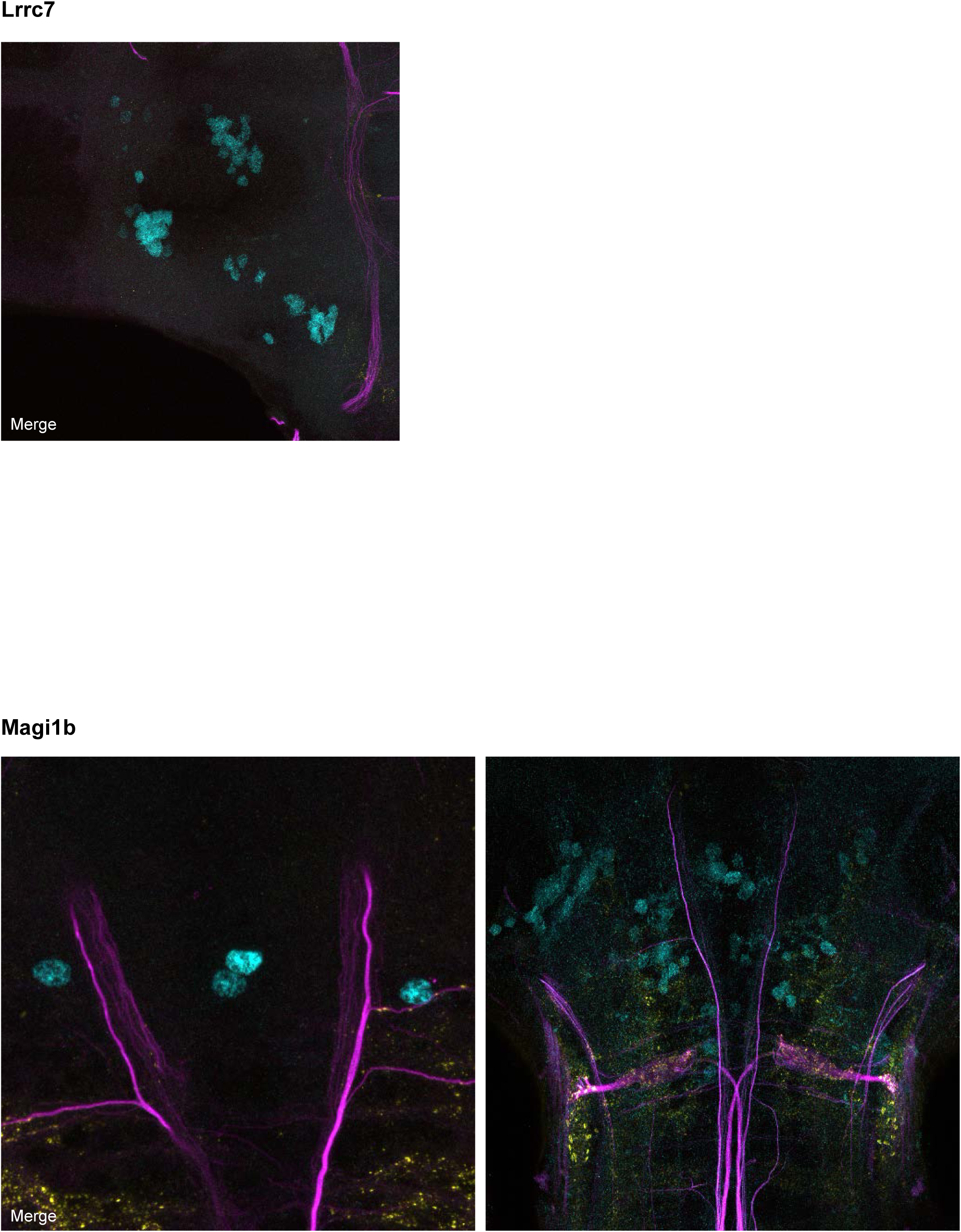
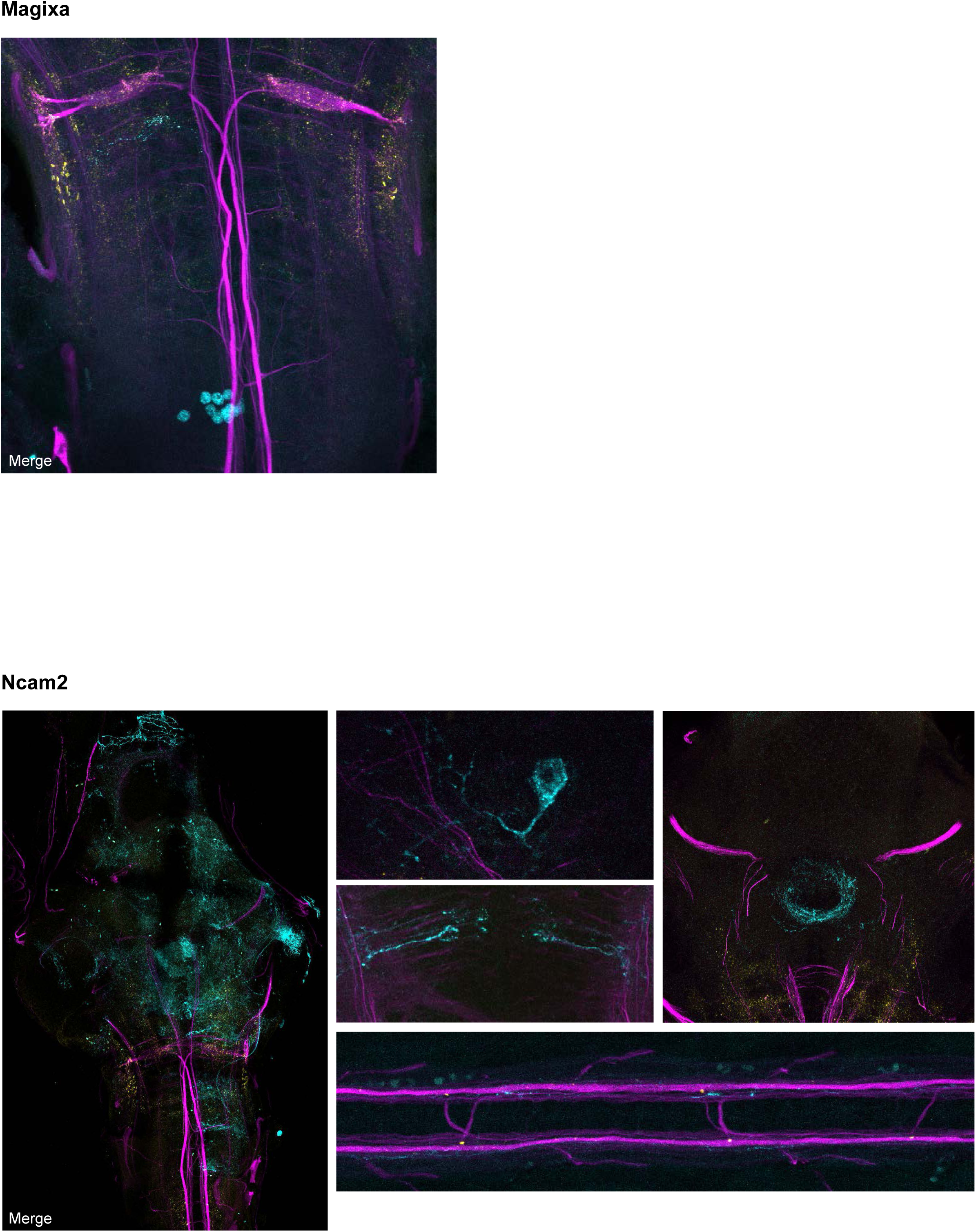
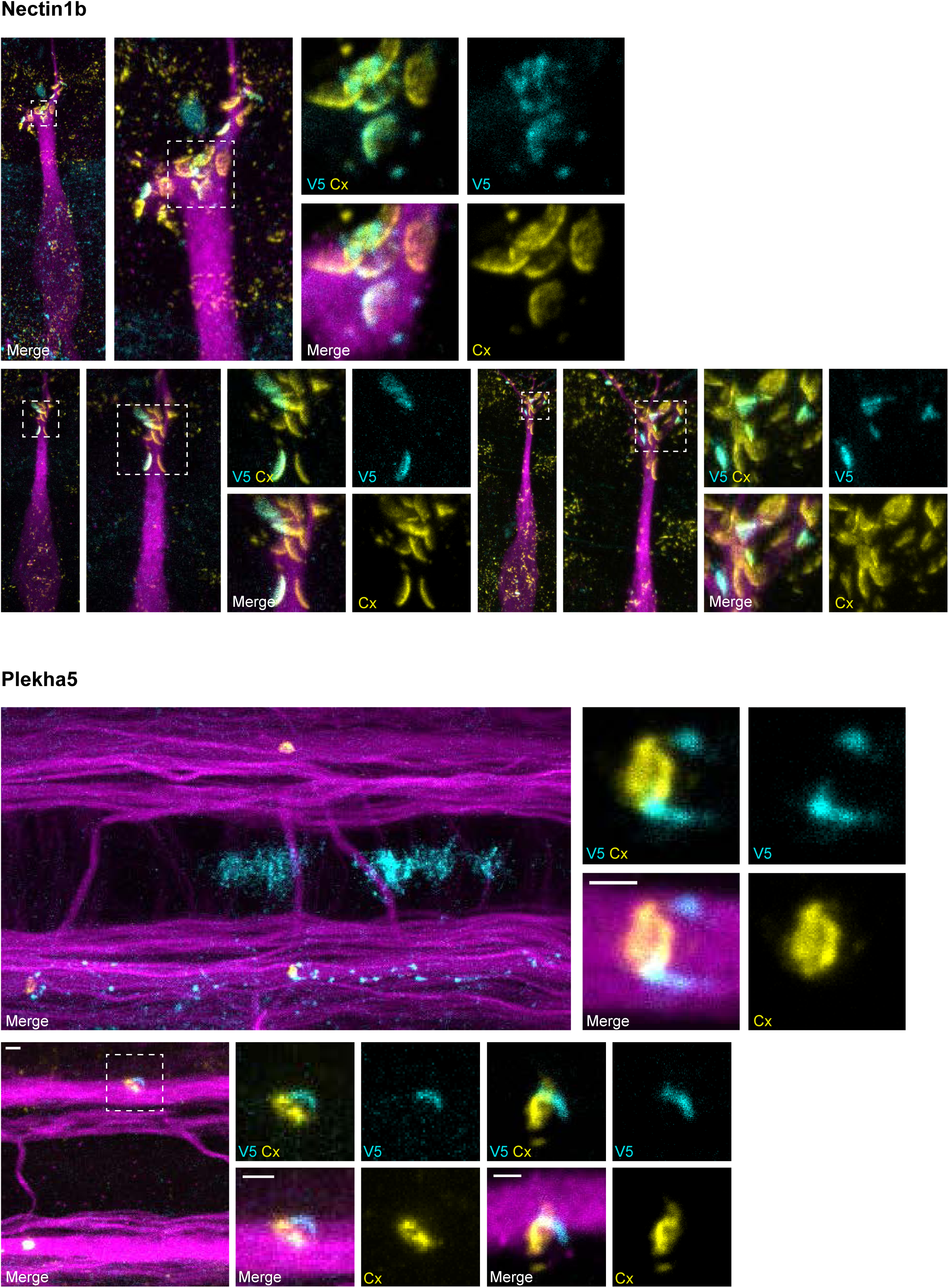
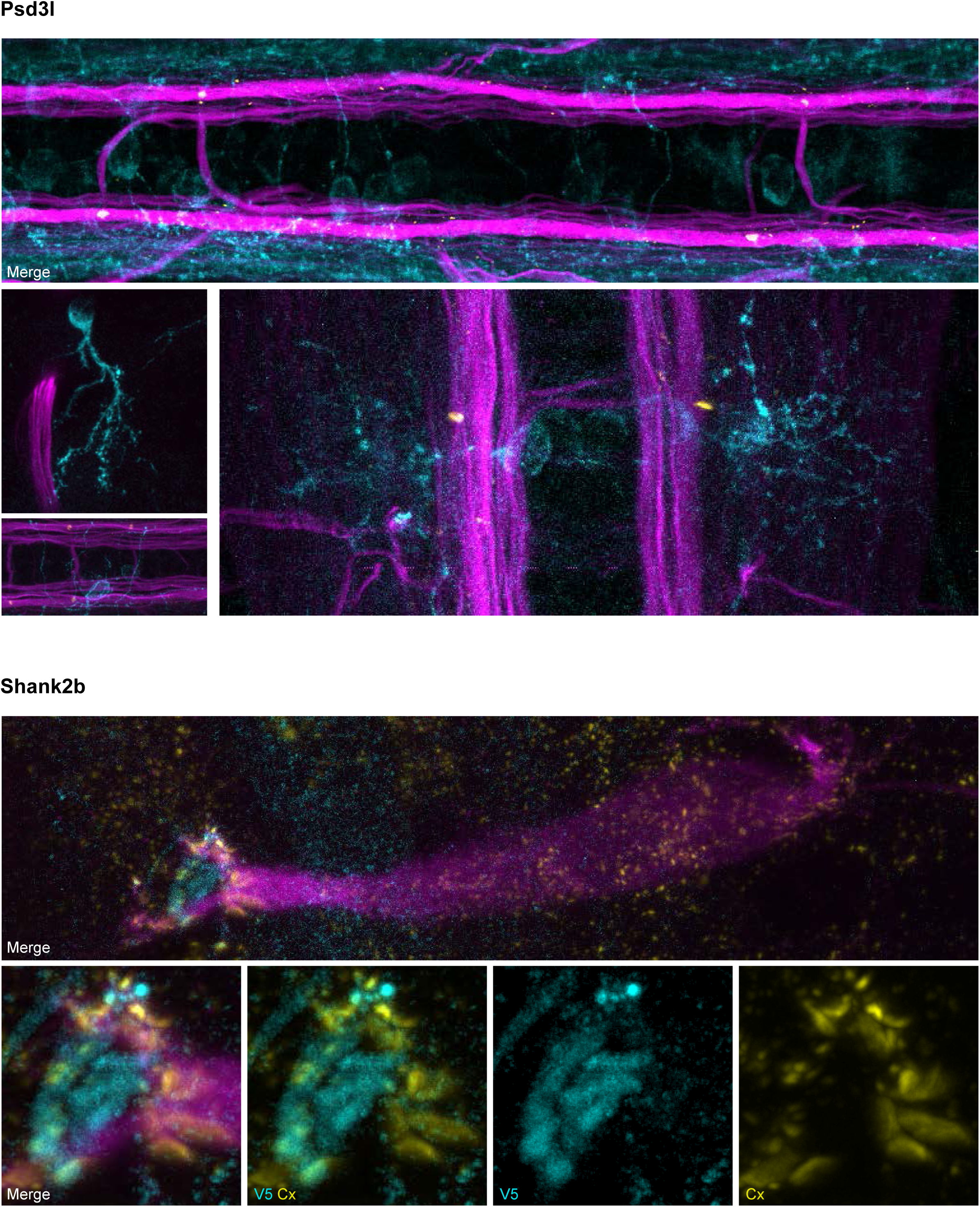
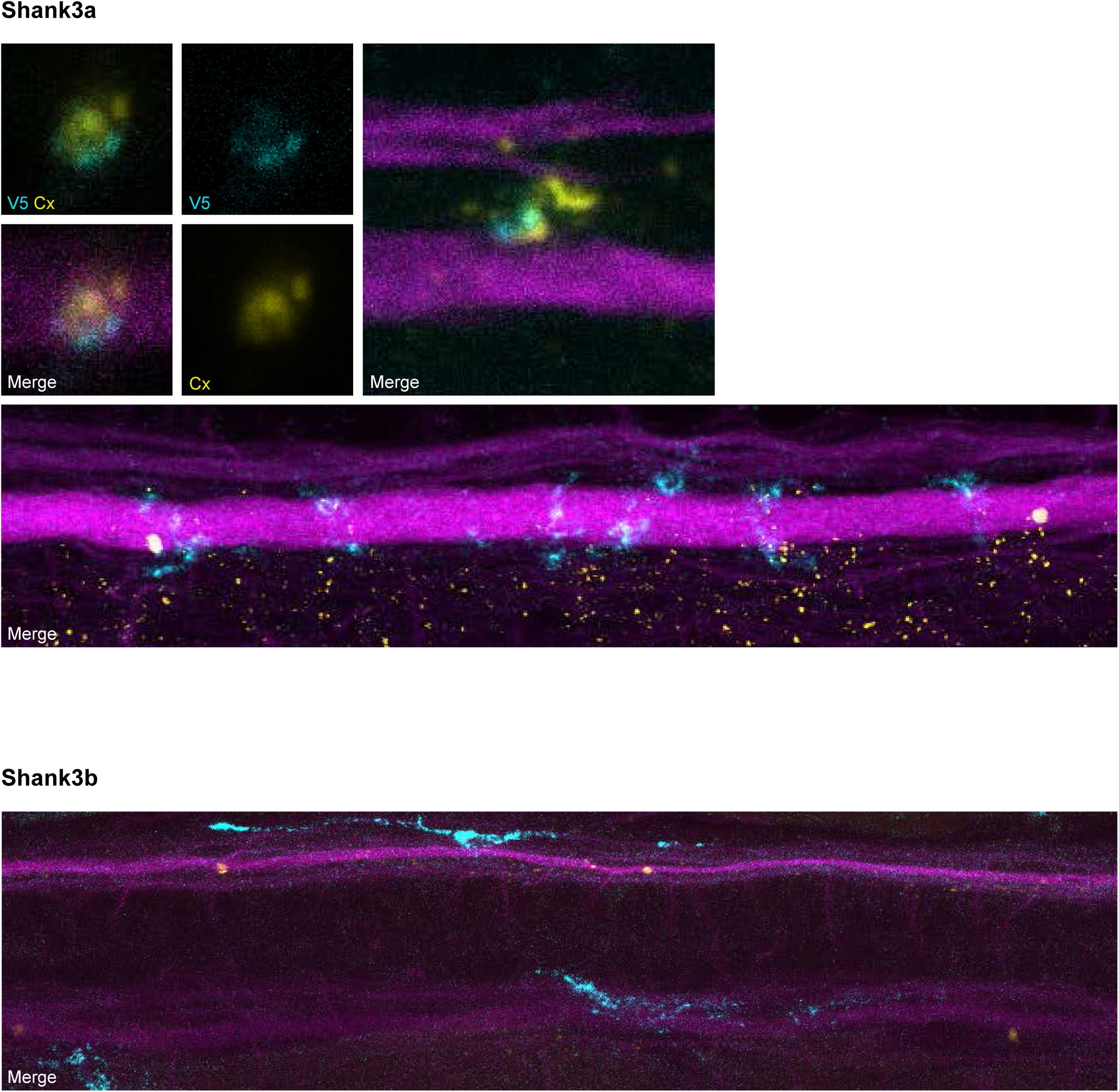
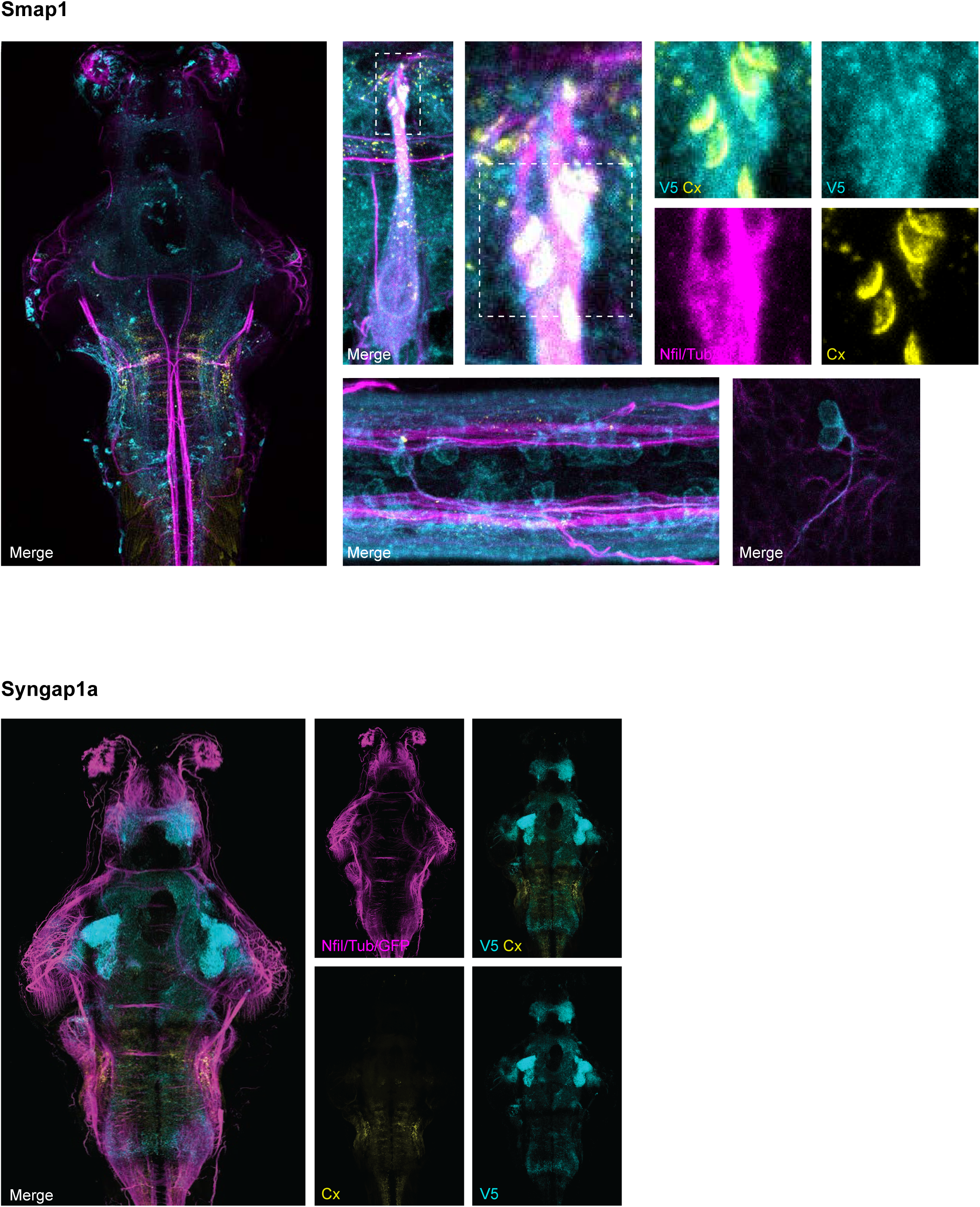
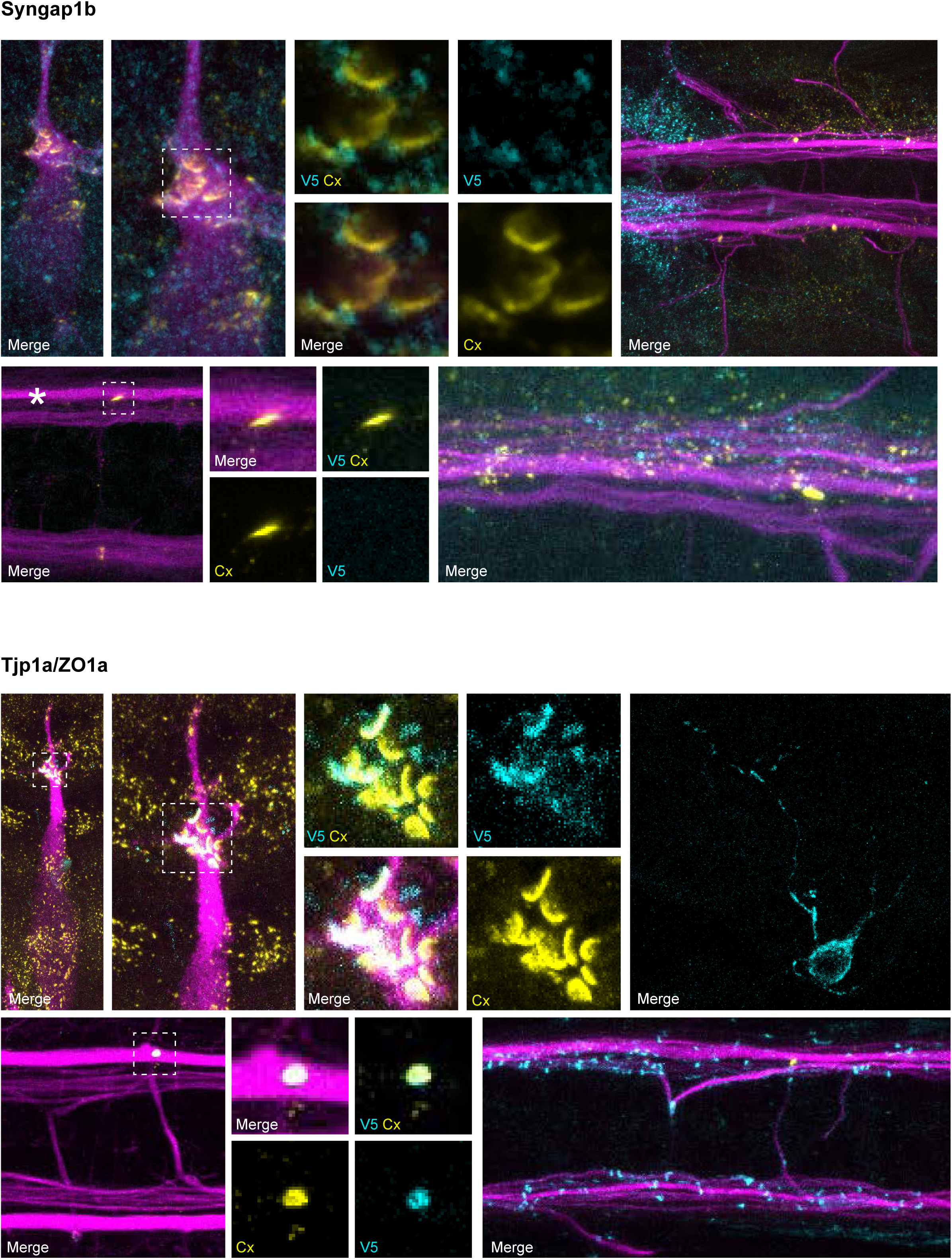
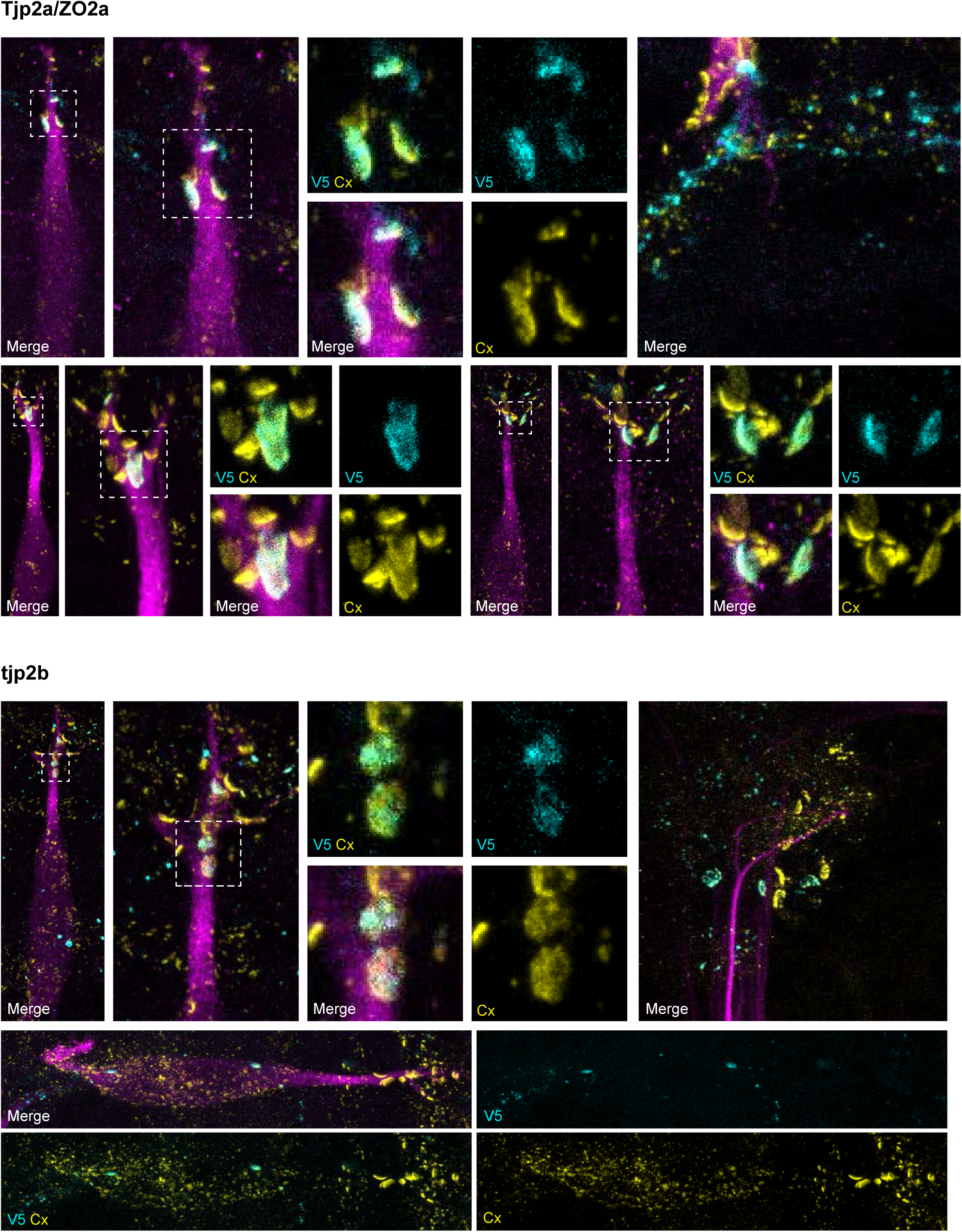
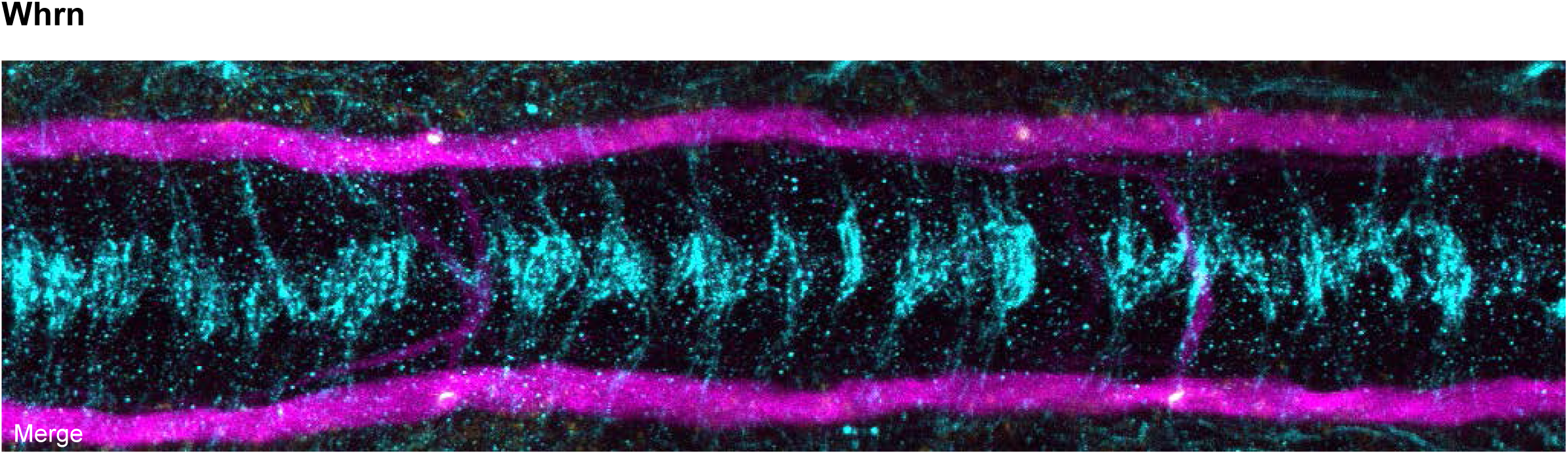
Mosaic V5-gene verification of candidate genes in developing zebrafish. Additional confocal images of V5-tagged candidate genes in 5-day-post-fertilization, *Et(T2KHG)^zf206^* zebrafish larvae. Genes are alphabetically ordered. Animals are stained with anti-GFP/anti-NF/anti-tubulin (magenta), anti-Cx34.1 (yellow), and anti-V5 (cyan). Neighboring panels show individual and merged channels, as labeled. Images are mainly tile scans of the whole brain or focused on hindbrain and spinal cord circuitry related to the M-cell, with examples from additional brain regions.

**Figure S6.**
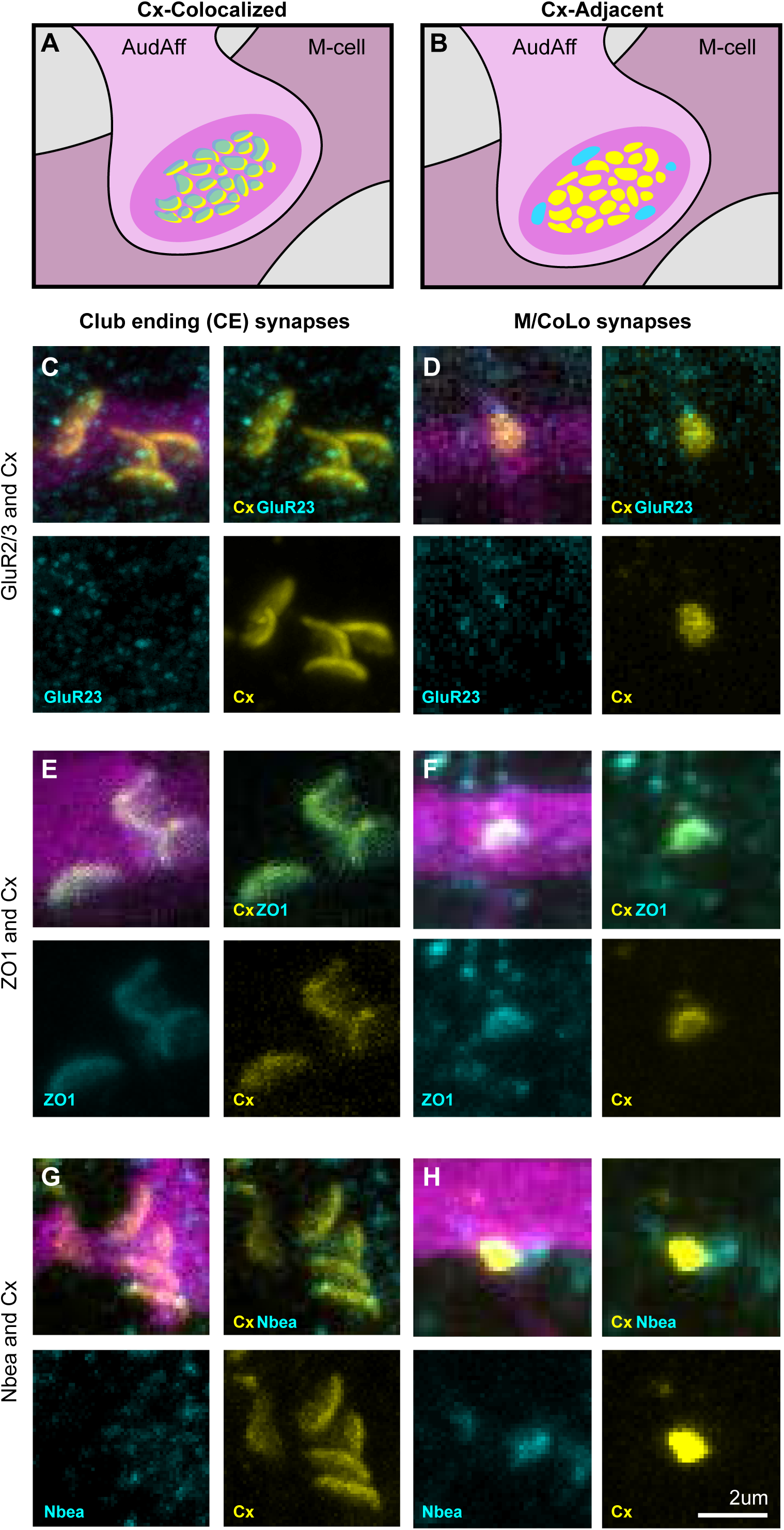
Connexin-colocalized and connexin-adjacent models of electrical synapses. **(A,B)** Model of the M-cell CE mixed electrical/chemical synapses illustrating Connexin-colocalized (A) or Connexin-adjacent (B) patterns. Gap junction plaques made of Connexin protein are yellow, while synapse-localizing proteins are shown in cyan. **(C-H)** Confocal images of Connexin-associated genes in the Mauthner circuit in 5-day-post-fertilization, *Et(T2KHG)^zf206^* zebrafish larvae. Animals are stained with anti-GFP/anti-NF/anti-tubulin (magenta), anti-Cx34.1 (yellow), and anti-GluR2/3 (cyan, C,D), anti-ZO1 (cyan, E,F), or anti-Nbea (cyan, G,H). Scale bar = 2 µm.

